# Circuit specific specialization of human basal ganglia astrocytes

**DOI:** 10.64898/2025.12.19.695583

**Authors:** Yuanyuan Fu, Nelson J. Johansen, Niklas Kempynck, Wubin Ding, Meghan A. Turner, Aaron D. Garcia, Matthew T. Schmitz, Jennie Close, Inkar Kapen, Madeleine Hewitt, Stephanie C. Seeman, Brian Long, Song-Lin Ding, Windy Ho, JT Mahoney, John K. Mich, Boaz P. Levi, Amit Klein, Maria Margarita Behrens, Joseph Ecker, Stein Aerts, Rebecca D. Hodge, Ed S. Lein, Trygve E. Bakken

**Affiliations:** Allen Institute for Brain Science; Seattle, WA, 98109; Laboratory of Computational Biology, VIB Center for AI & Computational Biology, Leuven, Belgium; VIB-KU Leuven Center for Brain & Disease Research, Leuven, Belgium; Department of Human Genetics, KU Leuven, Leuven, Belgium; Genomic Analysis Laboratory, The Salk Institute for Biological Studies, La Jolla, CA 92037

**Keywords:** basal ganglia, striatum, glial heterogeneity, astrocyte-synapse interactions, single-nucleus multiomics, chromatin accessibility, DNA methylation, enhancer-AAV

## Abstract

Astrocytes shape synapses and circuits, yet human basal ganglia astrocyte diversity is incompletely defined. We built a multimodal atlas by integrating single-nucleus RNA-sequencing and chromatin accessibility with DNA methylation, 3D chromatin conformation, and spatial transcriptomics, then mapped basal ganglia programs onto a whole-brain reference. Astrocytes segregated into three anatomical subgroups spanning striatal gray matter, extra-striatal gray matter, and white matter, with subgroup-biased neurotransmitter transporters and synapse-associated programs consistent with differences in dominant afferent input. Within striatum, dorsal and ventral astrocyte populations aligned with distinct microcircuits and were conserved in nonhuman primates. A deep learning sequence model identified subgroup-associated enhancer code and, when benchmarked against published enhancer-AAV datasets, supported the design of candidate viral tools to target basal ganglia astrocyte programs *in vivo*. Together, these data define major axes of human astrocyte specialization and provide a framework for cell type-specific dissection of basal ganglia function.

## INTRODUCTION

Astrocytes are an abundant glial cell type in the adult brain and play essential roles in maintaining homeostasis, supporting neural function, and preserving the integrity of the blood-brain barrier.^1^ Far from being passive bystanders, astrocytes actively participate in synaptic formation, pruning, maturation, and neurotransmitter cycling.^2–5^ While traditionally categorized into white- and gray-matter astrocytes based on morphology and location, growing evidence indicates that this binary framework oversimplifies the molecular and regional diversity of astrocytes.^6–9^ Canonical markers such as *GFAP* and *TNC*, which are commonly used in the cortex to distinguish fibrous from protoplasmic astrocytes, ^7,10^ are broadly expressed across both gray- and white-matter structures in non-telencephalic (non-TE) regions^7^, thereby blurring marker-based distinctions between these astrocyte identities in subcortical structures.

The basal ganglia (BG) consist of evolutionarily conserved nuclei that collectively regulate motor control, cognition, and motivation.^11,12^ These interconnected nuclei include the caudate nucleus (Ca), putamen (Pu), and nucleus accumbens (NAC), which together form the striatum (STR), along with the external and internal globus pallidus (GPe and GPi), ventral pallidus (VeP), subthalamic nucleus (STH), and substantia nigra (SN). In rodents, striatal astrocytes have also been shown to influence local circuit dynamics and behavioral flexibility,^13–17^ providing functional evidence for circuit-specialized astrocyte states within BG pathways. This anatomical and functional complexity makes the BG an ideal system to explore the diversity of astrocytes in the context of defined input-output architecture.

Recent large-scale mouse^6^ and human^7^ whole-brain atlases have identified astrocytes in the STR that are molecularly distinct from astrocytes in other TE and non-TE regions. In contrast, astrocytes in the globus pallidus (GP) are more transcriptomically similar to astrocytes in the midbrain or thalamus than to those in the STR,^7^ raising fundamental questions about how astrocyte identity is defined within subcortical gray matter (GM) and to what extent astrocyte diversity in the BG reflects local anatomy, long-range connectivity, or developmental origin. A BG-focused, multimodal framework is therefore needed to place these whole-brain observations into a circuit- and region-resolved context.

Here, we delineate the molecular and spatial organization of astrocytes in the adult human BG using an integrated single-nucleus multiomic framework spanning transcriptomics, epigenomics, and chromatin architecture, together with spatial transcriptomic profiling, and then place these BG-defined programs into a broader whole-brain context. This analysis revealed three molecularly distinct astrocyte subgroups with different spatial localization across BG nuclei. Subgroup-specific gene programs link astrocyte molecular specialization to circuit context and synaptic input type, and many of these molecular features are shared beyond the BG. A deep learning sequence model decodes subgroup-specific enhancer code that can be leveraged to design BG astrocyte-targeting tools. Finally, we identified dorsal-ventral heterogeneity within the STR, pointing to locally specialized astrocyte states. Together, these findings redefine astrocyte diversity in the human BG and provide new insights into region-specific glial specializations in complex subcortical circuits.

## RESULTS

We investigated the molecular and regional diversity of human BG astrocytes by integrating multiple single-nucleus molecular datasets—including paired transcriptomic and chromatin accessibility profiles (10X Multiome),^18^ spatial transcriptomics (MERSCOPE),^19^ and joint chromatin conformation/DNA methylation profiling (snm3C-seq)^20^—that are described in companion articles spanning comparable BG regions (Figure 1A).

**Figure 1.**
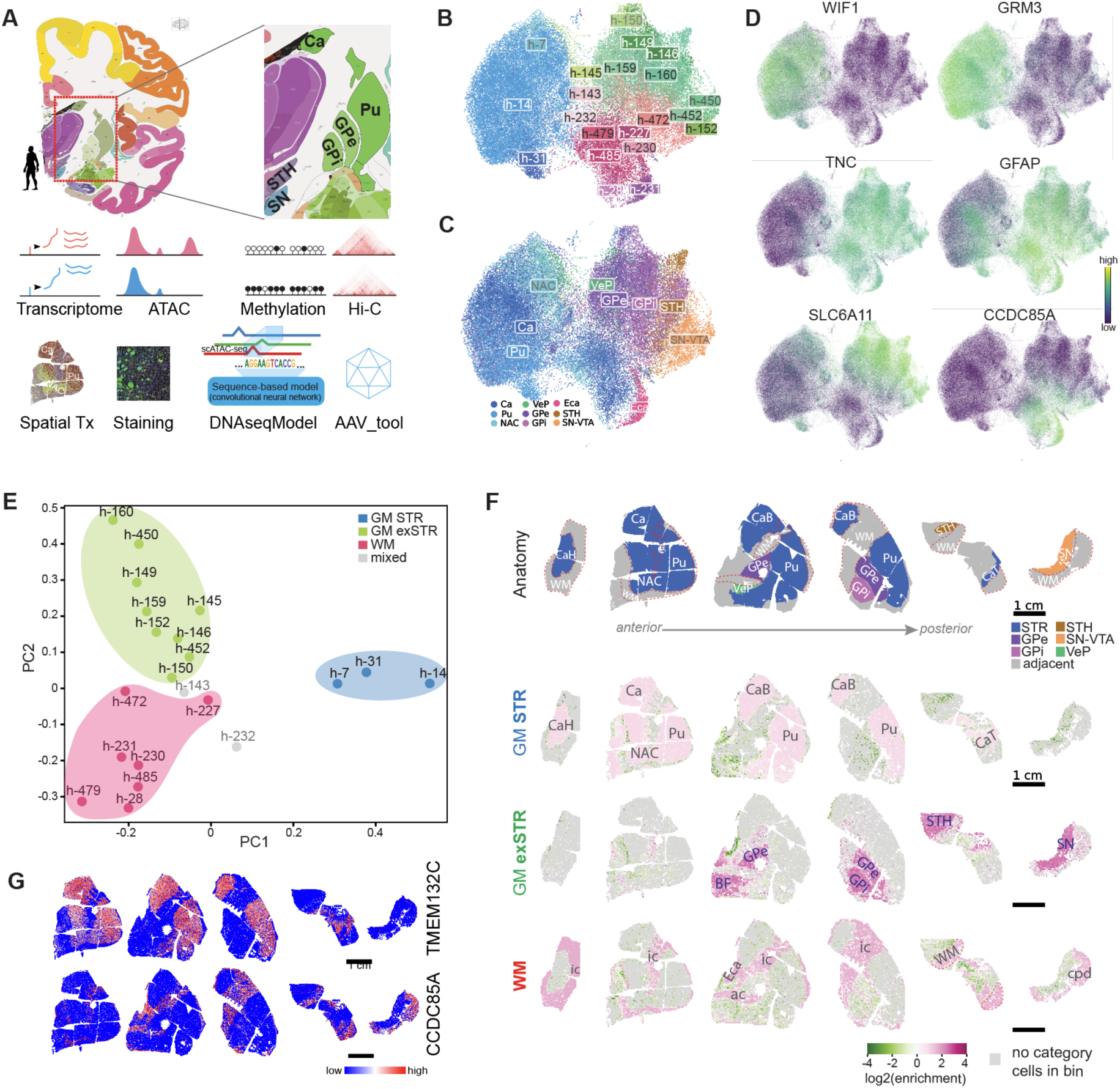
Regional specialization and spatial distribution of human BG astrocytes. **(A)** Schematic of sampling regions and multiomics datasets used in this study. **(B)** UMAP plot of human snRNA-seq astrocytes, showing cells colored by clusters. **(C)** UMAP plot of human snRNA-seq astrocytes, showing cells colored by sampling regions. **(D)** UMAP plots of human snRNA-seq astrocytes colored by well-known cortical astrocyte marker genes. **(E)** PCA plot of human BG spatial transcriptomics data (PC1 vs PC2). **(F)** Spatial enrichment maps of 3 human BG astrocyte subgroups across the BG tissue sections. Top row: BG nuclei are labeled, and white matter regions are indicated by dashed red lines. Bottom rows: Color scale corresponds to subgroup spatial enrichment, calculated as log_2_ of the ratio between the proportion of cells from the subgroup within each 300-µm grid cell (“local proportion”) and the proportion of cells from the same subgroup across that section (“global proportion”). Abbreviations: CaH, head of caudate; CaB, body of caudate; CaT, tail of caudate; BF, basal forebrain; ic, internal capsule; ac, anterior commissure; Eca, peri-caudate ependymal and subependymal zone; cpd, cerebral peduncle (crus cerebri). **(G)** Spatial expression maps of representative genes from the human BG MERSCOPE panel demonstrating clear patterns consistent with astrocyte subgroup distributions.

### Single-nucleus RNA-seq identifies transcriptional heterogeneity of human BG astrocytes

We first analyzed snRNA-seq data from 136,684 high-quality nuclei from 10 human donors (Figure S1) that were classified as astrocytes in a companion manuscript.^18^ Unsupervised clustering identified 21 astrocyte clusters that exhibited clear regional enrichment (Figures 1B and 1C). For example, clusters h-7, h-14, and h-31 were predominantly derived from the striatal dissections (Ca, Pu, and NAC), whereas clusters h-159 and h-160 were enriched in non-striatal regions (GP, STH, and SN; Figures 1B, 1C and S1F). To relate these clusters to known cortical astrocyte subtypes, we examined the expression of cortical GM astrocyte markers *WIF1* and *GRM3*, and cortical white matter (WM) markers *TNC* and *GFAP*.^7^ In a low-dimensional embedding of BG astrocytes, *WIF1* and *TNC* marked complementary domains (Figure 1D). Within the *TNC*-enriched domain, expression of the astrocytic GABA transporter gene *SLC6A11*^21,22^ and the cell adhesion related gene *CCDC85A*^23^ highlighted finer heterogeneity within this population (Figure 1D).

### Spatial transcriptomics divides astrocyte clusters into three anatomical subgroups

To determine the spatial distribution of the 21 snRNA-seq clusters, we analyzed human BG spatial transcriptomic data,^19^ where each MERSCOPE cell had been mapped to the best-match snRNA-seq cluster based on marker gene expression correlation. Consistent with the snRNA-seq patterns, principal component analysis (PCA) of cluster-level spatial composition—rather than gene expression—readily separated the *WIF1*-high, STR-enriched astrocyte clusters (e.g. h-14) from other clusters along PC1, while PC2 distinguished the *SLC6A11*-high (e.g., h-160) and *CCDC85A*-high (e.g., h-479) populations (Figure 1E). Density plots of individual clusters further revealed clear anatomical enrichment for 19 of the 21 clusters: h-7, h-14, and h-31 were enriched in GM STR; h-145, h-146, h-149, h-150, h-152, h-159, h-160, h-450, and h-452 in GM outside the STR (GM exSTR); and h-28, h-227, h-230, h-231, h-472, h-479, and h-485 in WM (Figure S2). Pooling the clusters assigned to each subgroup (GM STR, GM exSTR, and WM) produced density plots with strikingly clear spatial patterns (Figure 1F), mirroring both the PCA separation and the marker-defined UMAP distributions from the snRNA-seq data (Figures 1B-1E). Two small clusters (h-143 and h-232) displayed mixed gray- and white-matter spatial patterns and expressed marker genes from both compartments (Figures 1D and S2). We therefore classified them as ‘mixed’ groups, which may represent doublets that escaped stringent QC thresholds (Figures S1A and S1B). Consistent with these transcriptomic and spatial patterns, joint immunohistochemistry (IHC) and RNAscope labeling further supported the three astrocyte subgroups: *GRM3* expression was prominent in GM STR astrocytes, *SLC6A11* was elevated in GM exSTR, and GFAP fibers were substantially more abundant in WM and GM exSTR than in STR (Figure S3). Subsequent analyses focused on the three major astrocyte subgroups.

Examination of expression patterns in the spatial transcriptomic gene panels further highlighted candidate markers with clear spatial patterns: *TMEM132C*, *OAF*, *SEL1L3*, and *EPHB1* for GM STR astrocytes, and *CCDC85A*, *SMOC1*, *CD44*, and *VCAN* for WM astrocytes (Figures 1G and S4). In addition to subgroup specific markers, we identified genes with shared expression patterns across subgroups: *GPC5* was highly expressed in both GM STR and GM exSTR astrocytes, *RGS6* and *GRIK3* in GM exSTR and WM, and *DPP10* in GM STR and WM (Figure S4). These patterns were overall consistent with the expression profiles observed in the snRNA-seq modality (Figure S5A), supporting their robustness across platforms. Notably, many of these genes also exhibited broad expression in neuronal and immature oligodendrocyte populations within the full BG taxonomy (Figure S5B), indicating that while useful for distinguishing astrocyte subgroups, they are not astrocyte-restricted markers.

### Subgroup-specific transcriptional programs

Differential expression analysis of the snRNA-seq dataset delineated the molecular programs distinguishing the three astrocyte subgroups (Table S1). GM STR astrocytes showed the strongest subgroup-specific signatures, while GM exSTR astrocytes had few unique markers and shared many expression changes with WM (Figures 2A, 2B). WM astrocytes additionally exhibited distinct molecular features (Figures 2A, 2B). Several transcription factors (TFs) showed subgroup-biased expression: *FOXG1*, *EMX2*, and *LHX2* were enriched in GM STR astrocytes (Figure 2B), consistent with a TE program, whereas *SMAD9* and *CREB5* were more highly expressed in WM, and *TSHZ2* showed preferential expression in GM exSTR and WM (Figure 2B). Notably, *LHX2* and *CREB5* not only exhibited subgroup-biased expression but also resurfaced in our ATAC-based sequencing modeling, described below.

**Figure 2.**
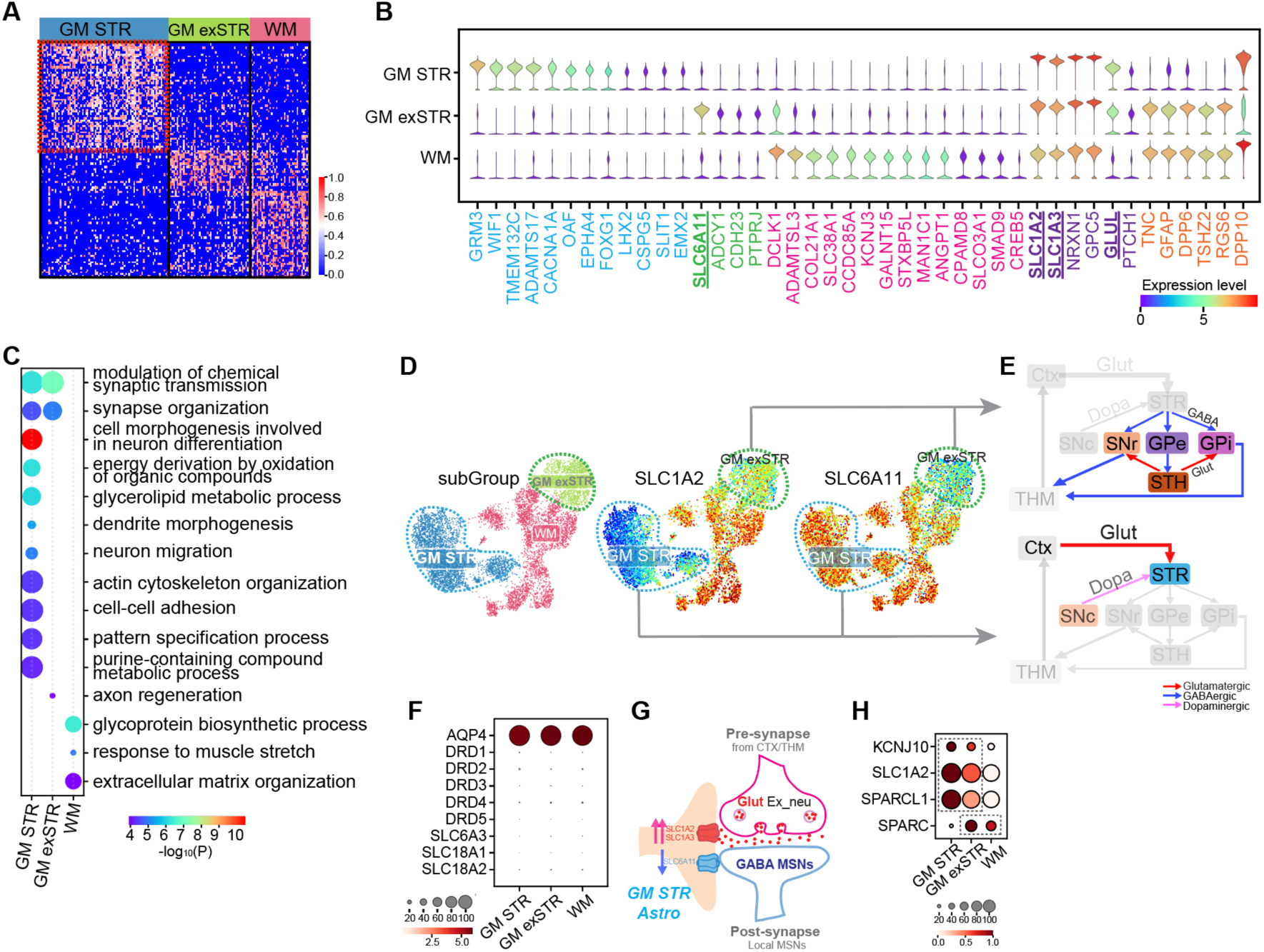
Molecular Signatures of 3 Human BG Astrocyte subgroups. **(A)** Heatmap displaying the expression patterns of subgroup-specific top DEGs across the astrocyte subgroups. **(B)** Violin plots illustrating the expression of the top differentially expressed genes identified for each of the three subgroups. **(C)** Gene Ontology (GO) enrichment analysis for genes enriched in GM STR, GM exSTR and WM astrocytes. Each dot represents an enriched biological process. Dot color, enrichment significance (-log_10_ P value). Dot size, gene set size of GO terms. **(D)** snm3C-seq methylation profiles of *SLC1A2* and *SLC6A11*. Genes encoding major neurotransmitter transporters and enzymes critical for neurotransmitter uptake show divergent patterns across the three astrocyte subgroups. **(E)** Simplified schematics of BG circuitry. The diagram depicts the major connections between cortex (Ctx), striatum (STR), substantia nigra pars compacta (SNc), substantia nigra pars reticulata (SNr), external and internal segments of the globus pallidus (GPe and GPi), subthalamic nucleus (STH), and thalamus (THM). Arrow colors indicate neurotransmitter types: red = glutamatergic (excitatory) input, blue = GABAergic (inhibitory) input, and pink = dopaminergic modulation. **(F)** Dot plot showing robust expression of a canonical astrocyte marker (AQP4) and very low expression of dopamine receptors and transporters in human BG astrocytes. **(G)** Schematic illustrating that GM STR astrocytes express higher levels of the glutamatergic transporter gene SLC1A2 rather than the GABAergic transporter gene SLC6A11, even though local neurons in the striatum are predominantly GABAergic. **(H)** Dot plot showing astrocyte-related genes involved in astrocyte-neuron interaction categories, illustrating their divergent expression patterns across the three subgroups.

WM astrocytes expressed extracellular matrix (ECM)-related genes such as *COL21A1*, *ADAMTSL3*, and *ANGPT1* (Figure 2B), consistent with Gene Ontology (GO) enrichment for ECM organization and glycoprotein biosynthesis (Figure 2C). These signatures underscore their structural and supportive roles of astrocytes in WM tracts. By contrast, both GM subgroups (GM STR and GM exSTR) were enriched for synapse-related terms, including modulation of chemical synaptic transmission and synapse organization (Figure 2C). We also noted that many of the top subgroup-biased differential expression genes (DEGs) showed broad neuronal expression rather than astrocyte specificity (Figure S6), consistent with their association with synaptic and circuit-related pathways.

### Synapse-interaction gene biases in astrocyte subgroups reflect circuit adaptation

In line with these functional enrichments, many synapse-related genes displayed subgroup-biased expression, emphasizing astrocyte-synapse interactions. As a key example, astrocytes at tripartite synapses use transporters to clear glutamate or GABA and the enzyme GLUL to convert them into glutamine for neuronal reuse.^21^ Consistent with this program, *GLUL* was elevated in both GM subgroups. The synaptic glutamate reuptake transporter *SLC1A2* (GLT1) was higher in GM astrocytes and most highly expressed in GM STR, whereas the GABA transporter *SLC6A11* (GAT3) was selectively enriched in GM exSTR but barely detectable in GM STR (Figure 2B), a bias also mirrored by DNA methylation profiles (Figures 2D and S7).

Although more than 90% of striatal neurons are GABAergic medium spiny neurons (MSNs), GM STR astrocytes preferentially expressed the glutamate rather than the GABA transporter. This asymmetry is consistent with circuit organization, as striatal territories are densely innervated by cortico-striatal excitatory inputs onto MSNs (Figure 2E), aligning with the elevated expression of *SLC1A2* in GM STR astrocytes. In contrast, GM exSTR astrocytes—sampled largely from GPe, GPi, STH, and SN—encounter dense GABAergic inputs from striatal MSN projections and GPe-STH connections, consistent with selective enrichment of *SLC6A11*. These patterns are consistent with astrocytic transporter expression aligning with the dominant neurotransmitter inputs of each region. We also examined dopamine receptors and transporters, given the extra dopaminergic innervation of the STR, but these were almost absent across astrocyte subgroups (Figures 2F and S8), suggesting that astrocytic specialization in the BG is more tightly coupled to glutamatergic and GABAergic circuits than to dopaminergic modulation (Figure 2G).

Beyond neurotransmitter cycling, additional functional modules reinforced this specialization. The inwardly rectifying potassium channel *KCNJ10* (Kir4.1),^24^ well known for coupling with glutamate transporters to maintain extracellular K⁺ and glutamate homeostasis^25,26^, mirrored the GM STR-biased expression of *SLC1A2* (Figure 2H). Similarly, the synaptogenic factor *SPARCL1* (Hevin) was enriched in GM STR, consistent with its role in bridging presynaptic neurexins and postsynaptic neuroligins to promote excitatory synapse formation and maturation,^27,28^ whereas its antagonist *SPARC*,^2,28^ was lowest in GM STR and conversely higher in GM exSTR (Figure 2H). Consistent with these transcriptional differences, DNA methylation patterns for both genes (Figure S7I; Table S2) closely paralleled their RNA profiles, further supporting the subgroup-specific regulatory landscape revealed in GM STR and GM exSTR.

### Deep learning sequence model reveals subgroup-specific gene regulatory logic

To investigate the gene regulatory mechanisms driving the specialized expression of astrocyte subgroups, we trained a deep learning model (CREsted^29^) to predict chromatin accessibility per subgroup from genomic sequences (Figure 3A). By learning transcription factor binding motifs and their combinatorial logic underlying subgroup-specific accessibility, the model provides an interpretable link between sequence features and biased accessibility, offering insight into the structure and logic of the underlying gene regulatory networks (GRNs). It showed a significant correlation between predicted and observed accessibility profiles (Pearson’s r=0.40, P<0.001; Figure S9A) and maintained high performance when evaluated within each subgroup (Figure 3B).

**Figure 3.**
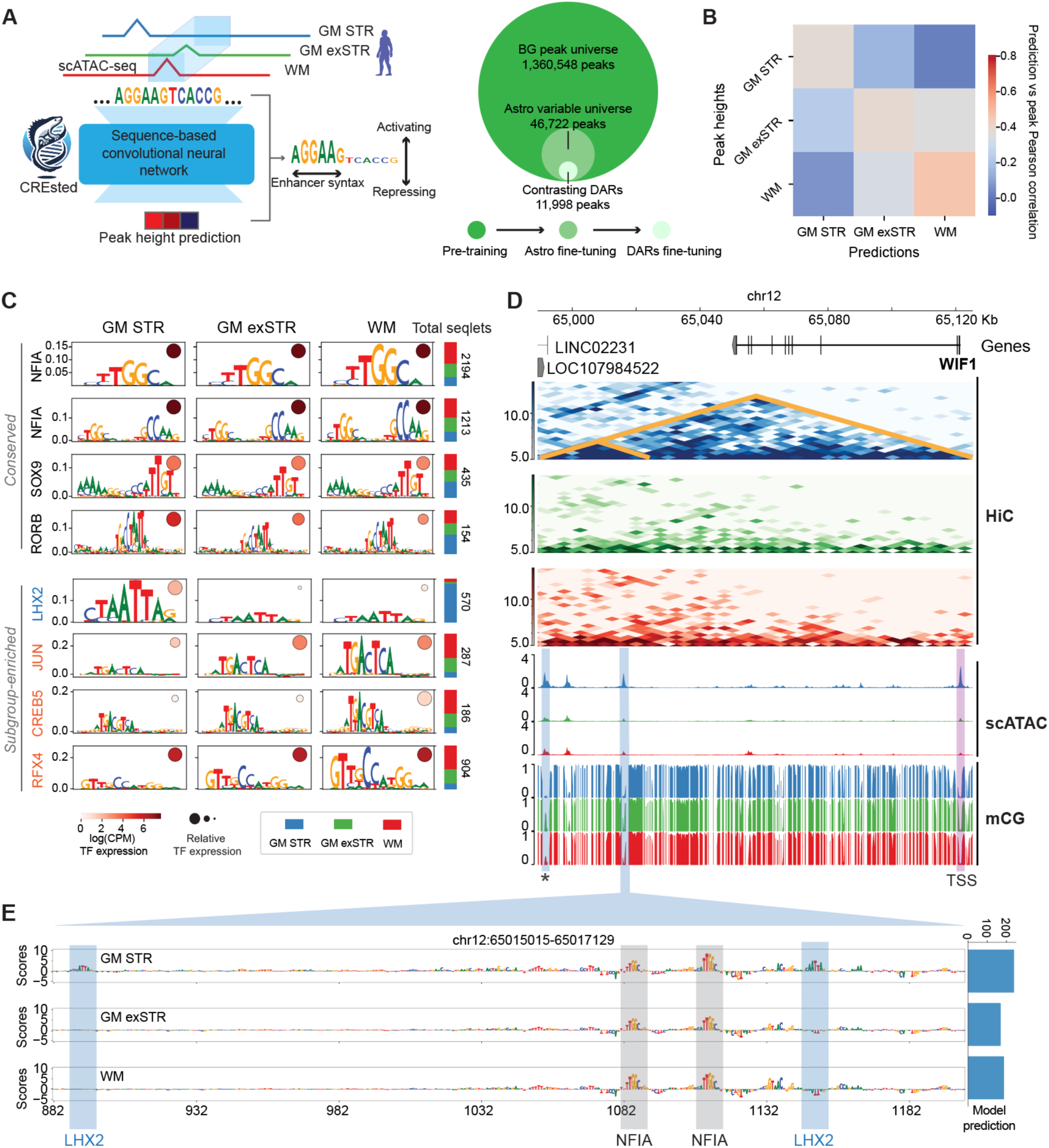
Deep learning model reveals regulatory motifs reinforced by multiomics. **(A)** Schematic illustration of sequence model (CREsted) training. Trained a CREsted model first on the whole human BG peak universe, then fine-tuned to specific astrocyte regions, then fine-tuned to DARs of human BG astrocyte subgroups. The DeepHumanAstroBG model predicts the aggregate peak height over the three subgroups. **(B)** Overview of the Pearson correlations between the log-transformed predictions and peak heights per subgroup to show the model performance. **(C)** Motifs identified by the model in the top 500 most specifically accessible regions for each subgroup. Motif heights are the average contribution scores from all instances identified across all subgroups. The distribution of motif instances found in each subgroup region set is shown with bar plots on the right. The absolute TF expression (logCPM) of proposed motif-TF matches is shown in color. Relative TF expression (per row divided by maximum value) is indicated by dot size. **(D)** Multiomic view of the *WIF1* gene locus, with Hi-C contact maps, and snATAC-seq and mCG (methylated CG dinucleotides) tracks. **(E)** The sequence model highlights a GM STR-specific regulatory region inside the *WIF1* gene locus harboring two LHX2 motifs and predicts this region to be selectively accessible in GM STR astrocytes.

The model identified the most frequently occurring TF motifs within subgroup-specific regions of open chromatin. Motifs for well-established astrocytic regulators, including NFIA-like, SOX9-like, and RORB-like motifs,^30–33^ were shared across all subgroups, and the corresponding TFs were highly expressed (Figure 3C). In contrast, LHX2 emerged as a driver of GM STR-specific peak accessibility, whereas JUN-like, CREB5, and RFX4-like motifs predominated in non-GM STR regions, consistent with expression of these TFs (Figure 3C). Finally, enrichment analysis of subgroup-specific differentially methylated regions in the snm3C-seq dataset^20^ further supported the predictions of our model, identifying, for example, LHX2 motif enrichment and its hypomethylation in GM STR astrocytes (Figure S9C). The model did not resolve distinct enhancer patterns between GM exSTR and WM astrocytes (Figure 3C), consistent with their broadly shared transcriptional, methylation, and Hi-C profiles despite their spatial segregation (Figures 1C-1F, 2A-2B, S2, S4, S5A, and S7).

To reveal potential subtype-specific regulatory mechanisms, we integrated model predictions with multiomic data for the *WIF1* locus, a GM STR marker relative to all other human BG cell populations (Figure S9D). There was concordant evidence for GM STR-specific *WIF1* regulation across data modalities: Hi-C (top panels) identified a topologically associating domain (TAD) encompassing the *WIF1* promoter, gene body and distal regulatory elements; snATAC-seq confirmed increased accessibility at the promoter and these distal sites; and snm3C-seq demonstrated lower methylation levels at these regions in GM STR (Figure 3D). Using the model to interpret sequence-level contributions to chromatin accessibility, we found two LHX2 and two pan-astrocyte NFIA motifs in a putative enhancer for *WIF1* in GM STR (Figure 3E). The accessibility of this region was predicted to be astrocyte-specific across many human brain regions with a similar motif-importance profile (Figures S9E and S9F) using a published model^33^ trained with human TE snATAC-seq data.^34^ This contrasts with another distal region (asterisk in Figure 3D) that, although predicted to be enriched for LHX2 motifs in GM STR (Figure S9G), is not astrocyte-specific (Figure S9H) and therefore represents a less compelling candidate for GM STR-specific WIF1 regulation. Similarly, we examined the *SPARC* locus, a gene with down-regulated expression in GM STR, and found that a region proximal to the promoter exhibits reduced chromatin accessibility and increased methylation in GM STR, with the model predicting the presence of a JUN motif (Figure S10). Collectively, these findings show how integrating sequence model predictions with multiomic data can decode cell type regulatory logic, pinpointing candidate elements underlying key marker genes such as *WIF1*.

### TF grammar of enhancer-AAVs

Building on these regulatory insights, we next examined whether the model predicts *in vivo* enhancer function and reveals the sequence features underlying enhancer activity. To this end, we applied the model to a set of experimentally validated astrocyte enhancers from recently published AAV enhancer toolboxes.^35–37^ Predicted accessibility patterns closely matched viral labeling (Figures 4 and S11), supporting the model’s ability to anticipate functional enhancer activity. For example, AiE2120m was predicted to be more accessible and AiE2121m less accessible in GM STR, consistent with their in vivo labeling: AiE2120m marked GM STR astrocytes and AiE2121m labeled GP astrocytes (Figures 4A and 4B). Contribution maps further revealed the TF motifs shaping these activities, with LHX2 and RFX4 emerging as potential drivers (Figure 4C).

**Figure 4.**
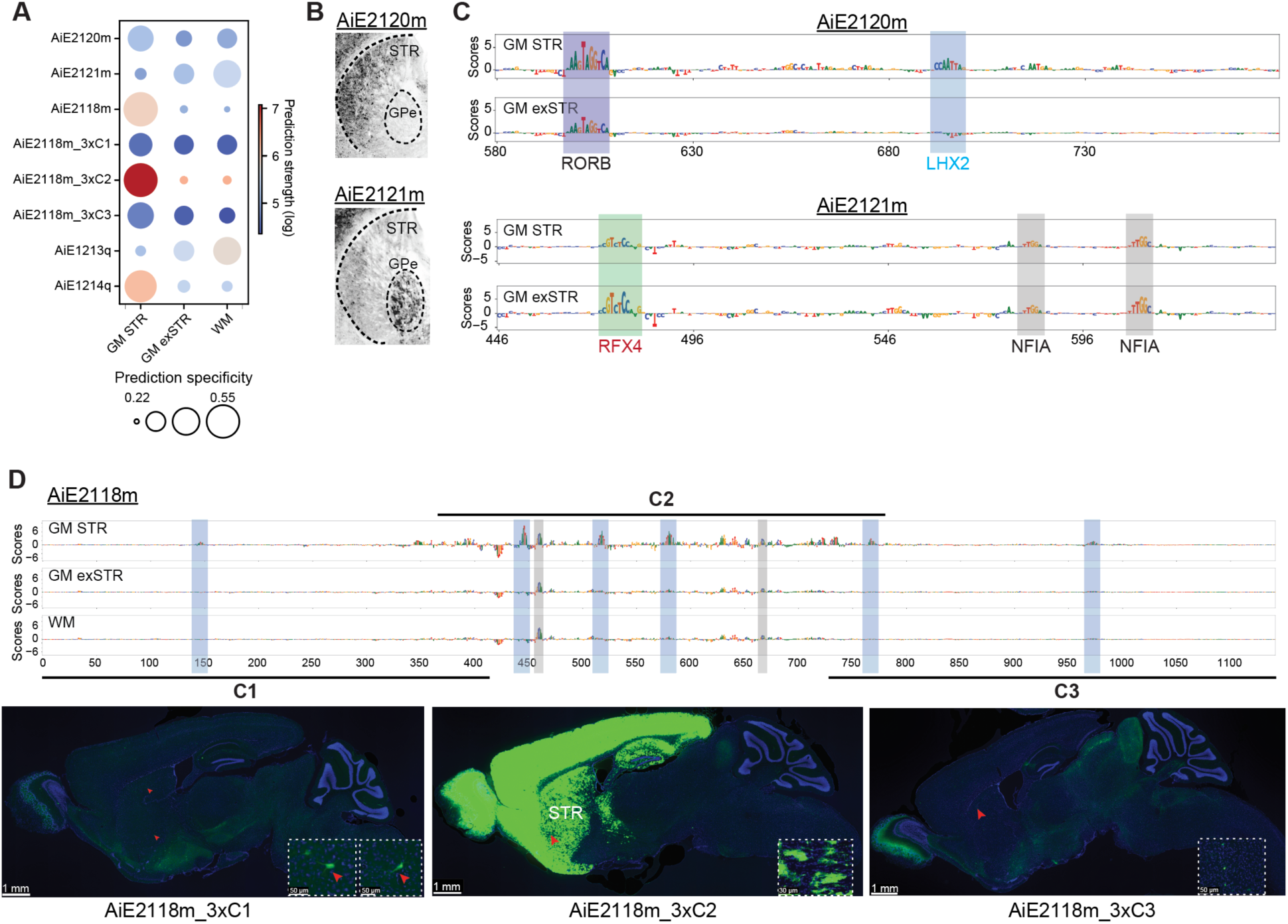
Sequence-based predictions of *in vivo* subgroup specific enhancer activity. **(A)** Model predictions of enhancer-AAV activity in BG astrocyte subgroups. Dot size reflects the specificity of each prediction, while color represents the log-transformed prediction score. **(B)** Predicted AAV enhancer activity from (A) matches the actual AAV labeling results from Mich et al. 2025.^35^ **(C)** Single-nucleotide view of base contribution scores in model-predicted AAV enhancers. The contribution scores of AiE2120m and AiE2121m give an idea on the underlying enhancer logic that allows for their specific expression pattern in the BG. The identified motifs follow the proposed candidates in Figure 3D: AiE2120m shows stronger LHX2 motif activity, consistent with GM STR, whereas AiE2121m shows stronger RFX4 motif activity, consistent with GM exSTR. **(D)** Example of a core-bashing AAV enhancer. The model predicted higher accessibility for C2, which was confirmed by much stronger viral labeling in STR.

We next asked whether model interpretability could predict functional activity across published enhancer fragments. For the AiE2118m enhancer, prior *in vivo* “core-bashing” experiments tested three fragments (C1–C3) derived from the original sequence, enabling segment-level assessment of enhancer function. The model assigned the central C2 fragment the highest contribution scores and predicted preferential accessibility in GM STR, driven by clustered LHX2 and NFIA motif instances consistent with astrocytic regulation. *In vivo* data agreed: C2 labeled GM STR astrocytes and adjacent TE astrocytes, whereas C1 and C3 produced only sparse labeling that appeared neuron-like by morphology (Figure 4D). Together, these results indicate that the model can predict functional differences among enhancer fragments and highlight the sequence features underlying subgroup-biased activity, providing a framework to prioritize candidate sequence edits for enhancer optimization.

### BG subgroup markers generalize to whole brain

To place the BG-derived astrocyte subgroups into a whole-brain context, we compared them with a recently published human brain atlas.^7^ This atlas delineates TE and non-TE astrocytes and GM and WM cortical astrocytes (Figures 5A and S12A). Label transfer mapping our three BG subgroups onto this reference revealed correspondence between GM STR and a striatal (TE) cluster (c55); GM exSTR and non-TE clusters enriched in thalamus, midbrain, and globus pallidus (c60 and c62); and WM and a TE WM cluster (c61) (Figure 5B). Thus, BG subgroups aligned with the principal subdivisions of astrocytes across the brain.

**Figure 5.**
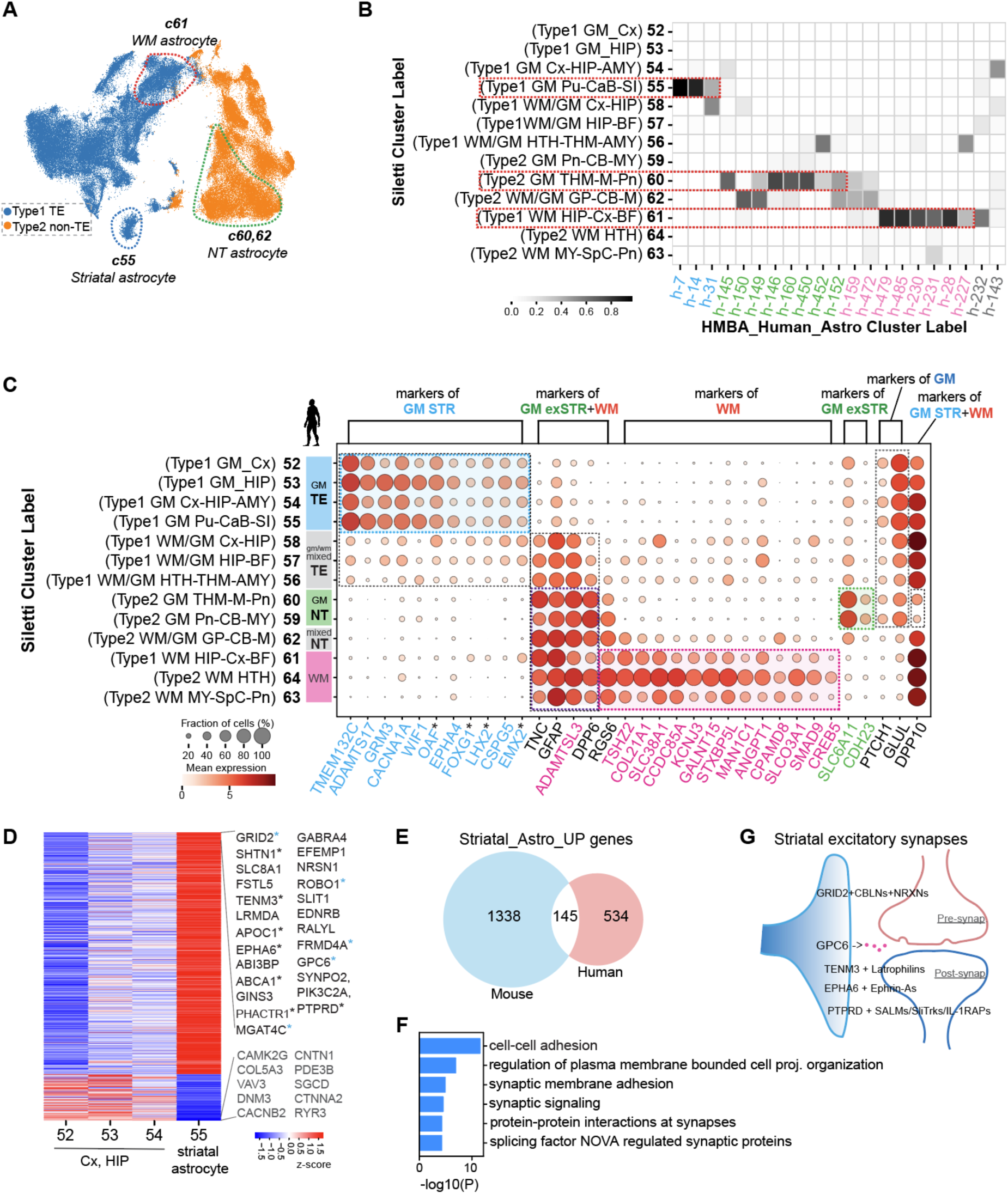
BG astrocytes share expression with major divisions of whole-brain astrocyte diversity. **(A)** UMAP plot of human astrocytes from the Siletti et al. whole-brain astrocyte taxonomy, showing telencephalic (blue, TE, Type 1) and non-telencephalic (orange, non-TE, Type2) compartments, with gray- and white-matter subdivisions outlined by dashed boxes. **(B)** Mapping of human BG astrocyte clusters onto the whole-brain reference. Color indicates the fraction of each BG cluster mapped to the corresponding whole-brain cluster (columns sum to 1). **(C)** Whole-brain expression patterns of BG subgroup marker genes identified in Figure 2B. **(D)** DEGs between human striatal and non-striatal telencephalic GM astrocytes from the Siletti et al. whole-brain taxonomy reveal a distinct transcriptional signature. Representative genes are indicated alongside the human heat map. Asterisks mark those also upregulated in mouse, with blue asterisks denoting genes ranked within the top 20 in both species. **(E)** Venn diagram showing that 145 genes are upregulated in both human and mouse striatal astrocytes. **(F)** GO enrichment analysis showing that 145 conserved upregulated genes are involved in synaptic adhesion and signaling. **(G)** Schematic illustrating how conserved top genes such as *GRID2* contribute to key components of the striatal tripartite excitatory synapse.

To identify features of BG subgroups that generalize to other brain regions, we examined the expression of the representative markers introduced in Figure 2. Strikingly, these genes stratified astrocytes across human brain, consistently separating TE from non-TE and distinguishing GM from WM compartments (Figure 5C). Many of these human markers were attenuated or lost in the mouse brain^6^ (Figure S12B). In contrast, the human GM STR marker *LHX2* was conserved in mouse and broadly expressed across TE astrocytes (Figures 5C and S12B), consistent with the functional activity in mouse of LHX2 motif-enriched enhancers (e.g., AiE2118m, Figure 4D), and sequence model predictions supporting activity across TE astrocytes (Figures S12C and S12D). Analysis of whole-brain developmental datasets^38^ showed that *LHX2* is highly expressed in TE astrocytes and their precursors in mouse, macaque, and human embryos (Figure S13), indicating that this molecular distinction is established during development and likely reflects differences in developmental programs and regional patterning.

Despite conservation of markers and key developmental TFs such as *LHX2*, the striatal astrocytes (c55) showed a distinct transcriptional signature compared with other TE astrocytes (Figures 5D and S12E; Table S3), and 145 genes were also upregulated in striatal astrocytes in mouse (Figure 5E; Table S4). These conserved genes were strongly enriched for synaptic adhesion and signaling functions (Figure 5F) and converge on excitatory synapse organization (Figures 5D and 5G). *GRID2*, which ranks among the top three markers in human and mouse, promotes excitatory synaptogenesis through cerebellin-neurexin interactions,^39,40^ while the astrocyte-secreted GPC6 supports excitatory synapse formation and stabilization^41^; adhesion-related factors such as ROBO1 further contribute to circuit refinement. Together, these analyses extend the striatal astrocyte-synapse signature identified in Figure 2 beyond the BG context, indicating that this excitatory synaptic program is a recurring feature of TE striatal astrocytes relative to other TE astrocyte populations.

### Local heterogeneity of GM STR astrocytes reveals dorsal-ventral differences

After establishing the molecular distinctiveness of striatal astrocytes, we next examined local heterogeneity within the GM STR subgroup. This subgroup comprises three transcriptionally defined clusters h-14, h-31, and h-7. Cluster h-14 represents the dominant population and is broadly distributed across the STR, whereas h-31 and h-7 are less abundant populations with more restricted anatomical patterns. Analysis of dissection proportions across donors revealed that h-31 is enriched in the dorsal STR (STRd), especially the Pu, while h-7 is localized predominantly to the NAC (Figure 6A). Although three donors in h-7 showed a higher proportion from Ca rather than NAC, closer inspection revealed that these Ca samples were from the CaH (Figure 6A), which lies very close to the NAC and may reflect minor mis-dissections and/or a rostral–caudal gradient in h-7 enrichment. These regional sampling patterns are consistent with cellular density maps derived from spatial transcriptomic data (Figure S2).

**Figure 6.**
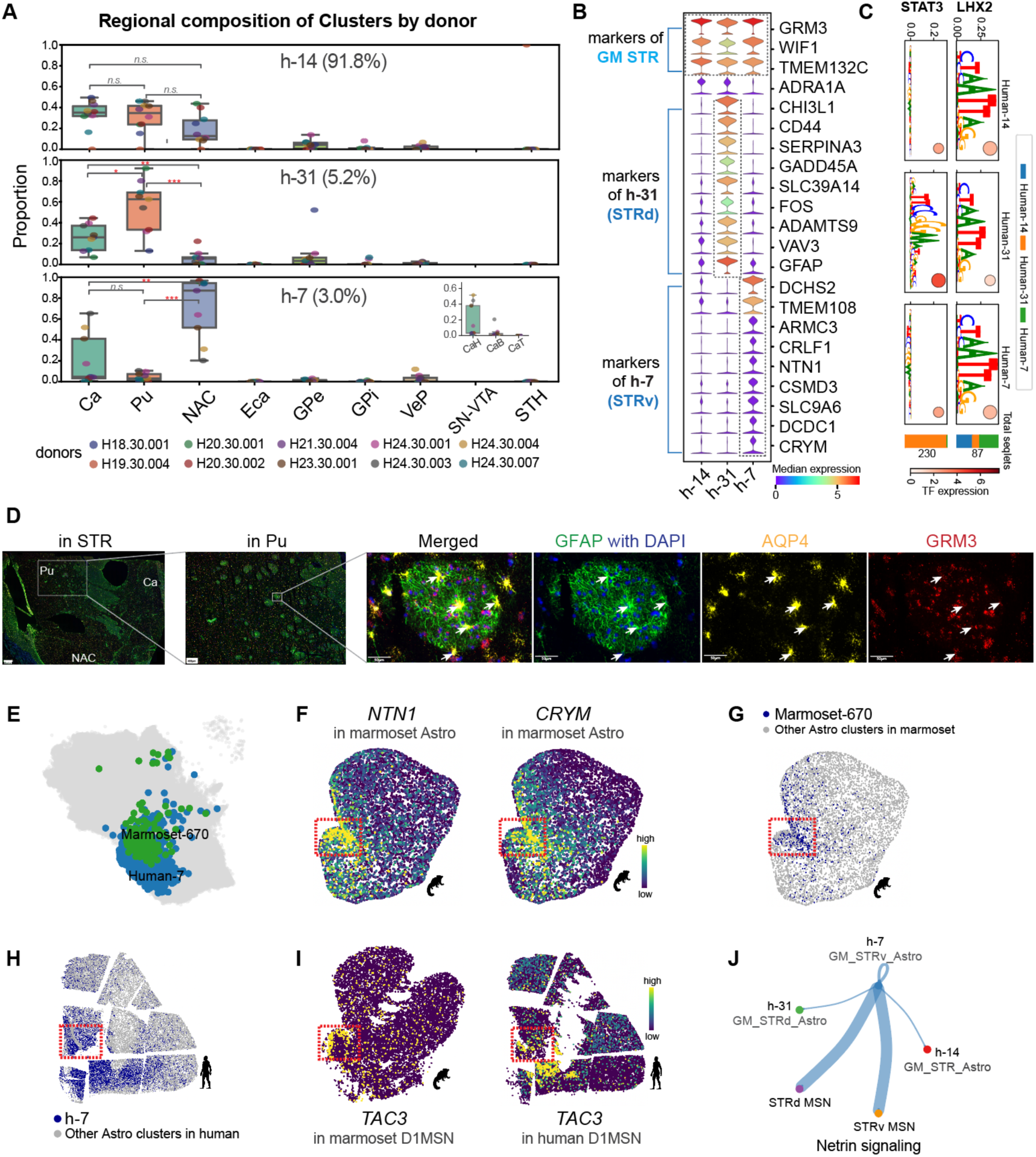
Local heterogeneity of GM STR astrocytes reveals dorsal-ventral differences. **(A)** Box plots showing the regional composition of clusters (h-14, h-31, and h-7) within the GM STR astrocyte subgroup across donors. Each point indicates the proportion of cells from a given region per donor (values normalized to 1 within each donor). Mann-Whitney U test; *, p < 0.05; **, p < 0.01; ***, p < 0.001; n.s., not significant. CaH, head of caudate. CaB, body of caudate. CaT, tail of caudate. **(B)** Candidate marker genes of clusters within GM STR astrocytes. **(C)** Motifs identified by the DeepHumanAstroSTR model in the top 500 most specifically accessible regions for each cluster. Motif heights are the average contribution scores from all instances identified across all GM STR astrocyte clusters. The distribution of motif instances found in each cluster region set is shown with bar plots on the right. The absolute TF expression (logCPM) of proposed motif-TF matches is shown in color. Relative TF expression (per row divided by maximum value) is indicated by dot size. **(D)** Joint IHC and RNAscope labeling for GFAP (green), DAPI (blue), AQP4 (yellow), and GRM3 (red) in the human striatum. Left: Low-magnification view showing STR anatomy, including Ca, Pu, and NAC. Center-left: Higher-magnification image of Pu highlighting GFAP⁺ WM tract patches. Center-right to right: Zoomed view of a representative patch and corresponding single channel images (Merged, GFAP with DAPI, AQP4, GRM3), illustrating co-expression of AQP4 and GRM3 in GFAP⁺ astrocytes (arrows). **(E)** UMAP plot of integrated human and marmoset astrocytes from the snRNA-seq datasets.^18^ The human h-7 cluster and the Marmoset-670 are highlighted. **(F)** Spatial transcriptomic maps of *NTN1* and *CRYM* expression in the marmoset Astrocyte Group, generated from the companion paper by Hewitt et al. **(G)** Spatial transcriptomic map of cluster Marmoset-670 and other astrocytes in marmoset Astrocyte Group. **(H)** Spatial transcriptomic map of cluster Human-7 (h-7) and other astrocytes in human Astrocyte Group. The red dashed box indicates the NACsmd region. **(I)** Spatial transcriptomic maps of *TAC3* expression in STR D1 MSN Subclass of marmoset (left) and human (right), with the NACsmd region outlined by a red dashed box. **(J)** Cell-cell communication analysis via CellChat2 suggesting h-7 as an active Netrin-signaling sender interacting with both ventral and dorsal MSNs.

Differential expression analysis revealed clear dorsal-ventral molecular differences among GM STR astrocytes (Table S5). The abundant h-14 population exhibited relatively few unique markers (Figure 6B; Table S5). By contrast, the dorsal-enriched cluster h-31 showed elevated expression of genes closely associated with reactive or stress-responsive astrocyte states,^42^ including *CHI3L1*, *CD44*, *SERPINA3*, *GADD45A*, *FOS*, and *GFAP* (Figure 6B). CREsted modeling of peak accessibility across the three astrocyte clusters identified STAT3, a putative reactive astrocyte marker,^42^ as the primary driver of h-31-specific accessibility, accompanied by strong enrichment of AP-1 motifs in the h-31 regulatory elements (Figures 6C and S14A). Together, these features point to a reactive, state-associated regulatory program rather than a distinct lineage-defining enhancer code. In comparison, the model did not detect a similarly robust enhancer signature separating h-7 from the abundant h-14 cluster (Figure S14A), suggesting more limited cis-regulatory differentiation at this level. Consistent with this, joint DNA methylation and chromatin conformation profiling did not reveal a clear dorsal-ventral segregation among GM STR astrocytes at this scale (Figure S7). Multiplex staining further showed that within the Pu, the region where h-31 is most enriched, numerous GFAP⁺ WM tracts course through the tissue, and astrocytes in these subregions are predominantly AQP4⁺GRM3⁺GFAP⁺ (Figure 6D). Cluster h-31 (AQP4⁺GRM3⁺GFAP⁺) astrocytes may therefore encounter distinct extracellular and structural conditions along these tract-associated gray-white matter interfaces, contributing to elevated expression of reactive genes.^43,44^

Differential expression analysis further revealed a distinct set of ventrally enriched genes defining the h-7 cluster (Figure 6B; Table S5). Among these, *DCHS2* encodes a calcium-dependent cell-adhesion molecule and *TMEM108* encodes a synaptic transmembrane protein, and both were broadly expressed across BG neurons with limited expression in MSNs (Figure S14B), including selective enrichment of *DCHS2* in the ventral STRv D1 MSN Group (Figure S14C). In contrast, *ARMC3* and *CRYM* are not broadly expressed across BG neurons yet show pronounced ventral bias within MSNs, particularly in STRv D1 and STRv D2 MSN Groups, mirroring their ventral astrocytic enrichment and suggesting coordinated ventral molecular specialization across neuronal and non-neuronal cell types. Notably, canonical MSN markers such as *DRD1* and *DRD2* are nearly absent from all GM STR astrocyte clusters (Figure S14E), excluding ambient RNA contamination as the source of these shared ventral signatures.

### Ventral CRYM^+^/NTN1⁺ GM STR astrocytes are enriched in the primate NAc-smd

We next asked whether the ventral h-7 population exhibits a similarly defined anatomical niche. Direct evaluation in human tissue is limited because key h-7 markers, including *NTN1* and *CRYM*, are absent from the human spatial gene panel and anterior NAC coverage is sparse. To address this gap, we leveraged the primate BG and the mouse whole-brain transcriptomic and spatial atlases,^6,18,19^ which together provide broader spatial and molecular context for striatal astrocytes. Among these datasets, only the human and marmoset atlases resolved multiple striatal astrocyte clusters, enabling further characterization of their local diversity. Marmoset-670 cluster aligns closely with human h-7 (Figure 6E) and expresses the markers *NTN1* and *CRYM* that show strong, selective expression in the mediodorsal subdivision of NAC shell (NACsmd) (Figures 6F, 6G, and S15C–15E). Human h-7 astrocytes show an enrichment within the NACsmd (Figure 6H), while also extending into broader regions of the NAC, indicating a conserved but not spatially exclusive ventral specialization across primates.

These findings raise the question of whether astrocyte specialization in the NACsmd reflects neuronal and circuit organization in that BG region. Recent work identified a novel TAC3-archetype among macaque D1 MSNs that is localized to the NAC medial shell and proposed that it may represent an additional corticostriatal pathway.^45^ We confirm that *TAC3* expression is highest in primate STRv D1 MSNs (Figures S15F and S15G). In marmoset, *TAC3*^+^ D1 MSNs show a restricted spatial distribution along the medial margin of the NAC shell, closely corresponding to the NACsmd (Figure 6I). In human, *TAC3* expression is similarly elevated in the NACsmd but also extends into additional shell and core of the NAC (Figure 6I), mirroring the broader NAC distribution observed for h-7 astrocytes (Figure 6H). A similar species-dependent pattern is observed for the *HTR7*⁺ subgroup of the novel STRd D2 StrioMat Hybrid MSN population identified in our companion study.^18^ In marmoset and macaque, *HTR7*⁺ D2 StrioMat Hybrid MSNs are largely confined to the NACsmd, whereas in human they are distributed more broadly across the ventral STR while remaining enriched in NACsmd-associated regions (Figure S3.3 in reference 18).

The convergent spatial localization of *NTN1*^+^ ventral astrocytes and distinct ventral MSN populations within the NACsmd suggests a potentially coordinated astrocyte-neuron specialization within this ventral microcircuit. In line with this spatial organization, cell-cell communication analysis identified activation of the Netrin signaling pathway, with *NTN1* acting as a prominent sender signal from h-7 astrocytes to MSNs in the human dataset (Figure 6J).

## DISCUSSION

In most brain regions, the diversity of local neuron types and interwoven nature of projection sources obscure whether astrocytic states reflect local or afferent influences. The STR, by contrast, offers rare clarity because its local neurons are overwhelmingly GABAergic, yet its major afferent synapses are glutamatergic. This separation between local and input transmitter identities, together with the well-defined circuit architecture of the BG, provides a useful set of regions for dissecting astrocyte organization.

Our multiomic and spatial analyses reveal three major astrocyte subgroups in the human BG and show that their molecular specialization is tightly linked to circuit context and developmental origin. Astrocyte subgroups showed graded differences in neurotransmitter cycling genes—*GLUL* elevated across GM, *SLC1A2* highest in GM STR and lowest in WM, and *SLC6A11* selectively enriched in GM exSTR—mirroring the dense glutamatergic input to GM STR and GABAergic projections to GM exSTR. Similar gradients extend beyond the BG where *SLC1A2* decreases from TE to non-TE GM and further to WM, while *SLC6A11* peaks in non-TE GM astrocytes.

Although both GM subgroups share functional signatures of active neurotransmitter recycling and neuron-astrocyte metabolic coupling typical of GM, GM exSTR astrocytes overall display molecular and regulatory features more closely resembling those of WM astrocytes. Across transcriptomic, methylation, chromatin accessibility, 3D chromatin interaction, and spatial dimensions, GM exSTR consistently aligned nearer to WM than to GM STR, with GM exSTR and WM sharing a highly similar motif signature. At the gene level, GM exSTR displayed few subgroup-specific markers. Both snRNA-seq and snmC-seq identified *SLC6A11* as one of the rare distinctive genes, while most other features were shared with adjacent subgroups—for example, *TNC* and *GFAP* with WM, and *GLUL* with GM STR. Consistent with these molecular similarities, regions containing GM exSTR astrocytes also displayed dense GFAP-positive fiber networks reminiscent of WM. Together, these findings indicate that GM exSTR represents a GM population that has partially adopted WM-like molecular programs. One possible contributing factor is that within the BG, extra-striatal nuclei contain fewer neurons and more heavily myelinated projection fibers than the STR^46^, which may create a local environment favoring WM-like astrocytic states. While GM exSTR astrocytes share many features with WM astrocytes, we identified WM-enriched markers with sharply delineated spatial patterns that more selectively label BG WM than classical markers such as *TNC* and *GFAP*, providing refined molecular tools for targeting BG WM astrocytes.

Within GM STR astrocytes, we identify substantial local heterogeneity. The most abundant cluster (h-14) is broadly distributed across STR, whereas two less abundant clusters are enriched in the dorsal STR (h-31) and the NAC in the ventral STR (h-7). Cluster h-31 displayed transcriptional and regulatory features reminiscent of a reactive-like astrocyte state, including elevated expression of stress-responsive genes and pronounced enrichment of STAT3 and AP-1 motifs. Staining showed that h-31 astrocytes are concentrated within GFAP⁺ WM tract patches in Pu, a gray-white matter interface microenvironment that may contribute to their reactive-like profile.^43,44^ Yet h-31 retains core GM STR features, including broad expression of GM STR markers, highly accessible LHX2-associated regulatory elements, and transcriptomic proximity to other GM STR clusters rather than GM exSTR or WM populations. These observations support the view that h-31 represents a GM STR astrocyte subgroup whose reactive-like signature likely reflects adaptation to a local niche, highlighting how fine-scale striatal architecture can shape astrocyte specialization.

In contrast, the *NTN1*⁺*CRYM*⁺ cluster h-7 marks a ventral astrocyte specialization anchored in the NACsmd across primates. While the niche is more spatially restricted in non-human primates, in human it extends into broader NAC territories, paralleling the wider distribution of specific ventral MSN archetypes and preserving a within-species correspondence between ventral astrocyte and neuronal specializations. This NACsmd receives specialized projections from the dorsal hippocampus^47^ and the anterior paraventricular thalamic nucleus,^48^ and cell-cell communication analysis further implicates h-7 derived NTN1 as a prominent signal to MSNs. Together, these observations suggest that this ventral astrocytic program may be shaped by both local and input circuit contexts within the BG.

Subregional specialization is also evident within GM exSTR and WM astrocytes. In GM exSTR, we identify BG nuclei-enriched clusters, and WM astrocytes are largely distributed along major BG WM tracts, with some clusters displaying focal regional biases, for example, h-159 in the GPe and h-231 near the peri-caudate ependymal and subependymal zones (Figure S2). Together with the GM STR findings and subregional diversification of MSNs,^18,19,45,49^ these patterns indicate that astrocytes specialize to reflect regional connectivity and local microenvironment.

Our data also highlight an axis of astrocyte specialization related to developmental lineage. Across BG astrocyte subgroups, *LHX2* emerged as a defining feature of the GM STR and a robust marker of TE astrocytes across species, supported by whole-brain transcriptomic, regulatory modeling, and enhancer data. Embryonic single-cell datasets in human, macaque, and mouse further revealed that *LHX2* is specifically enriched in TE astrocytes and their progenitors, alongside established TE regulators *FOXG1* and *EMX2*. However, across neural lineages, LHX2 shows a broader dorsal bias that extends into mesencephalon and diencephalon, in contrast to FOXG1, which remains largely TE-restricted. These patterns suggest that LHX2 marks TE astrocyte identity and may contribute to establishing or maintaining TE astrocyte programs, while also being reused in dorsal brain regions for neurons.^33,38,50^

The GP provides a notable exception to this astrocytic TE program. Although anatomically TE, GP astrocytes more closely resembled those in the STH and SN than neighboring striatal astrocytes, consistent with whole-brain datasets that grouped GP astrocytes within a “non-TE type 2” population, even though the GP is the only major TE structure represented in that type.^7^ *LHX2* expression was nearly absent in GP astrocytes, reinforcing their divergence from the canonical TE lineage. Beyond the lower neuron-to-glia ratio that may favor a WM-like astrocytic state for GP, developmental timing and origin likely also contribute: the GP is developmentally heterogeneous and emerges early in embryogenesis^51^ and in close spatial and temporal proximity to earlier-developing diencephalic and midbrain territories,^52^ potentially biasing its astrocyte lineage toward a non-canonical TE program. Together, these observations suggest that developmental history and regional context can blur traditional anatomical boundaries of TE, with *LHX2* providing a useful marker for TE astrocyte identity and the GP illustrating an exception to this rule.

Finally, our interpretable sequence model, benchmarked against published enhancer-AAV datasets, enables decoding of subgroup-specific regulatory logic and design of genetic tools for targeted astrocyte access. By integrating motif contribution scores with snATAC-seq and methylation profiles, it identifies regulatory elements whose predicted accessibility and motif content match subgroup-specific enhancer activity *in vivo*. The model recapitulates the astrocyte-targeting patterns of recent enhancer-AAVs, including differential labeling of GM STR versus GP astrocytes. This enables rational design of tools to access specific BG astrocyte populations and circuits.

In summary, we define astrocyte diversity in the human BG along intersecting, scale-dependent axes of transmitter cycling, GM versus WM-like programs, subregional microcircuit specialization, and developmental lineage. The STR illustrates that astrocyte states can be aligned with afferent transmitter identity; GM exSTR and WM show that GM location does not guarantee classical GM programs; and subregional heterogeneity underscores that astrocytes encode fine BG architecture alongside MSNs. *LHX2* and enhancer analyses provide an entry point into the developmental and regulatory mechanisms that couple astrocyte identity to brain-wide TE versus non-TE organization and offer tools for cell type- and circuit-specific perturbation in the BG.

### Limitations of the study

This study integrates transcriptomic, epigenomic, spatial, and histological data to define astrocyte diversity in the human BG, and the atlas is derived from a limited number of adult donors with uneven coverage across some structures and WM tracts. Very rare, highly localized, or border-zone astrocyte populations are therefore likely underrepresented, and additional donors, particularly from anatomically undersampled regions, will further refine the diversity described here.

In addition, links between astrocyte transcriptional programs, transmitter-defined inputs, and BG architecture are based on correlative analyses of postmortem tissue and integrated multiomic datasets. Functional studies, including targeted physiological recordings and cell type-specific perturbations will be required to determine how these molecular signatures translate into defined roles of astrocyte subtypes within striatal and extra-striatal circuits.

## RESOURCE AVAILABILITY

### Lead Contact

Further information and requests for resources and reagents should be directed to and will be fulfilled by the lead contact, Trygve E. Bakken.

### Materials Availability

This study did not generate new unique reagents.

### Data and Code Availability

- Sequencing data and enhancer validation data used in this project are all from previous studies and are publicly available. Accession numbers and links to the datasets are listed in the key resources table.
- All original code has been deposited at GitHub and is publicly available as of the date of publication. GitHub URLs and DOIs are listed in the key resources table.
- Any additional information required to reanalyze the data reported in this paper is available from the lead contact upon request.

## Supporting information

Supplemental_Figures_BG_Astrocyte

Table_S1_DEGs_filtered_aross_ASC_subgroups.xlsx

Table_S5_DEGs_filtered_across_GM-STR-3-clusters.xlsx

Table_S4_C5527-Mouse_upreg_1483genes_vsOtherTE.xlsx

Table_S3_C55-Human_upreg_679genes_vsOtherTE.xlsx

Table_S2_DMGs_Hypo_filtered_across_ASC_subgroup.xlsx

## ACKNOWLEDGMENTS

This publication was supported by and coordinated through the BRAIN Initiative Cell Atlas Network (BICAN) (https://braininitiative.nih.gov/research/tools-and-technologies-brain-cells-and-circuits/brain-initiative-cell-atlas-network). This work was funded by the Allen Institute for Brain Science and by the National Institutes of Health UM1MH130981 (ADG, BL, BPL, ESL, IK, JC, JKM, JTM, MAT, MH, MTS, NJJ, RDH, SCS, SD, TEB, WH, YF). Research Foundation Flanders (FWO) PhD fellowship 1SH6J24N & V428025N (NK). The authors thank the founder of the Allen Institute, Paul G. Allen, for his vision, encouragement and support.

## AUTHOR CONTRIBUTIONS

Tissue acquisition: ESL, RDH

Data generation: ADG, AK, ESL, JE, JTM, MMB, RDH, WD, WH

Spatial transcriptomic data generation: BL, JC, MH, MAT, SCS

Viral tool testing: BPL, JKM

Data analysis: ADG, BL, IK, JC, MH, MAT, MTS, NJJ, NK, SCS, TEB, WD, YF

Data interpretation: ADG, BL, BPL, ESL, IK, JC, JE, JKM, JTM, MAT, MH, MTS, NJJ, NK, RDH, SA, SCS, SD, TEB, WD, WH, YF

Writing manuscript: ESL, MTS, NK, RDH, TEB, YF

## DECLARATION OF GENERATIVE AI AND AI-ASSISTED TECHNOLOGIES

During the preparation of this work, the authors used OpenAI ChatGPT-5 to support certain coding tasks, assist with refining the main text to improve conciseness and readability. After using this tool, the authors reviewed and edited the content as needed and take full responsibility for the final content of the publication.

## STAR METHODS

### METHOD DETAILS

#### PCA Analysis on Spatial Transcriptomic Data

For each section, cells were binned on a 2D grid (300 µm), and the number of cells from each astrocyte cluster was counted within each bin. These per-bin counts were concatenated across sections to form a matrix of bins × clusters. Each row was normalized to proportions (sum = 1) and z-scored across bins to equalize feature scales; bins containing missing values were excluded. Principal component analysis (PCA) was then applied to this normalized matrix using scikit-learn. In the resulting PC1-PC2 scatterplot, each astrocyte cluster is represented by a point whose x and y coordinates correspond to its loadings on PC1 and PC2, respectively, derived directly from the PCA components matrix (components_[0, c], components_[1, c]).

#### Spatial Enrichment Analysis of Astrocyte Clusters and Subgroups

To quantify the spatial distribution of human astrocyte clusters across tissue sections, we selected QC-passed astrocytes (Group==“Astrocyte”) from the human spatial transcriptomic dataset^19^ and included only those derived from six representative coronal sections (z_order ∈ {3, 6, 9, 12, 15, 0}). The global proportion of each of the 21 clusters across all astrocytes was calculated and used as the expected baseline for enrichment analysis. Within each section, cells were binned on a two-dimensional spatial grid with 300 µm resolution, and the log₂ enrichment of each cluster was determined as the ratio between its observed frequency in each spatial bin and its global baseline. The resulting enrichment maps display one heatmap per section, highlighting regions where specific astrocyte clusters are locally over- or under-represented relative to their overall abundance. A similar procedure was applied to astrocyte subgroups, using their labels to compute global and section-level enrichment.

#### Differential Expression Analysis of snRNA-seq Data

DEGs across human BG astrocyte subgroups were identified using the *rank_genes_groups()* function in Scanpy^53^ with the Wilcoxon rank-sum test, Benjamini-Hochberg FDR correction, and “rest” as the reference group. Candidate marker genes reported in Table S1 were selected using the thresholds “pvals_adj< 0.01, logfoldchanges > 1, and pct_nz_group > 0.2” for each subgroup. Example genes shown in the figures represent the top candidates obtained under a more stringent version of this filtering scheme.

For differential expression analysis among the three GM STR astrocyte clusters (h-14, h-7, and h-31), we first performed donor balanced down sampling to mitigate the large differences in cluster sizes (with h-7 and h-31 being much smaller than h-14). We subsampled up to 100 cells per cluster-donor combination, and the resulting down sampled object was analyzed using the same *rank_genes_groups()* function described above.

#### Gene Ontology (GO) Enrichment Analysis

For GO enrichment analysis, we derived a high-confidence marker set for each astrocyte subgroup by applying a more stringent version of our DEG filtering criteria, including a lower adjusted P threshold (0.01) and a higher fold-change requirement (>1), along with more restrictive detection-rate cutoffs. These refined candidate marker genes of each subgroup were then submitted to Metascape^54^ for GO enrichment analysis under default settings. For visualization, we retained only highly significant “GO Biological Process” terms and excluded categories with very small or excessively large gene-set sizes to avoid unstable or overly broad annotations. The top enriched terms from each astrocyte subgroup were then merged into a unified term set and visualized as a single dot plot.

#### CellChat Analysis

To investigate potential ligand-receptor interactions between astrocyte clusters and dorsoventral MSN populations, we aggregated the fine-grained annotations into five major classes: h-14 (GM STR astrocyte), h-7 (GM STRv astrocyte), h-31 (GM STRd astrocyte), STRd MSN (STRd D1 Matrix, STRd D1 Striosome, STRd D2 Matrix, STRd D2 StrioMat Hybrid, STRd D2 Striosome), and STRv MSN (STRv D1, STRv D2, STRv D1 NUDAP). The MSN cells were extracted from the same human BG dataset described in the companion study.^18^ CellChat^55^ analysis was performed in R using the human ligand-receptor interaction database (*CellChatDB.human*). For signaling inference, we used the default CellChat settings with excluding the “Non-protein Signaling” category (*subsetDB(CellChatDB)*). Over-expressed ligands, receptors, and interaction pairs were identified using CellChat’s standard preprocessing workflow. The resulting cell-cell communication networks were visualized using CellChat’s pathway plots.

#### snm3C-seq data processing and analysis

snm3C-seq^56^, which jointly profiles DNA methylation and 3D chromatin interactions from the same nuclei, were obtained from our companion study^20^ and processed using the same computational workflow. Briefly, we computed 100-kb binned average mCG and mCH methylation profiles and performed PCA separately on each matrix. The resulting low-dimensional representations were concatenated and used to generate a joint UMAP embedding. To mitigate technical and biological batch effects arising from study of origin, sequencing technology, and donor, we applied Harmony^57^ correction prior to graph construction and Leiden clustering. Consensus cluster assignments were subsequently derived using the ALLCools^58^ (https://github.com/lhqing/ALLCools) *ConsensusClustering()* function.

For cell-type annotation, we assessed CG hypomethylation at canonical marker genes. Gene-level methylation was quantified as the normalized average methylation across the gene body with ±2 kb flanking regions, leveraging the established inverse relationship between neuronal gene-body methylation and gene expression in the brain.

Differentially methylated genes (DMGs) were identified by comparing normalized gene methylation between each cell type and all others using a Mann-Whitney U test with Benjamini-Hochberg correction. For each gene, we computed the median methylation difference and the ROC AUC to quantify discriminatory power. Hypo-DMGs were defined by AUC ≥ 0.8, adjusted *P* ≤ 0.05, and median Δmethylation < 0.

We then identified differentially methylated regions (DMRs) between different cell types (one versus rest) using ALLCools. To identify candidate transcription factor regulators associated with cell type-specific hypomethylated DMRs (hypo-DMRs), we extracted genomic sequences spanning -250 to +250 bp around center of all hypo-DMRs and scanned for transcription factor binding motif using Cluster-Buster^59^ to create database. Motif enrichment was then performed using pycistarget v1.1 (https://github.com/aertslab/pycistarget).^60^

For 3D genome analyses, single-cell Hi-C contact matrices generated by snm3C-seq were aggregated into cell type-resolved pseudobulk contact maps using scHiCluster^61^ (https://github.com/zhoujt1994/scHiCluster), based on the methylation-derived cell-type labels. Raw contact counts from the pseudobulk matrices were used for genome browser visualization.

#### Coverage Quantification and Peak Calling of snATAC-seq Data

Human BG astrocyte ATAC data were obtained from the 10X Multiome dataset described in the companion study^18^. Because each nucleus contains paired RNA and ATAC profiles, we directly used the RNA-derived astrocyte subGroup annotations to subset and retain the matched ATAC profiles. Bigwig coverage tracks for each human astrocyte subgroup were generated using the *export_coverage()* function in SnapATAC2^62^ with the hg38 reference genome. Coverage was calculated from ATAC insertion sites in 10-bp bins and ‘CPM’ normalized using ±100 bp windows around transcription start sites (TSSs). Peaks for each subgroup were called with the MACS3 (https://github.com/macs3-project) wrapper (*snapatac2.tl.macs3*) and merged across subgroups using *snapatac2.tl.merge_peaks()* to obtain the human BG astrocyte universe peak set used for CREsted model training.

The three GM STR clusters were processed using the same preprocessing workflow, including TSS-normalized bigwig generation and MACS3-based peak calling, prior to CREsted model training.

#### CREsted Enhancer Modeling for Human BG Astrocyte

We trained a CREsted^29^ model (DeepHumanAstroBG) on human astrocyte snATAC-seq data at the subgroup level. During data preprocessing, we normalized peak heights using the *crested.pp.normalize_peaks()* function, which estimates a normalization scalar for each subgroup based on the distribution of the most accessible (top 3%) peaks. This step ensures comparable accessibility scales across subgroups. The peak regression model was then trained following the default parameters. Consensus peaks were split by chromosome into training, validation, and test sets with chromosome 10 for validation, chromosome 18 for test, and the remaining chromosomes for training.

We first pre-trained our model on the whole human BG peak universe (1,360,548 peaks),^18^ ensuring that the model was trained on a large enough dataset to generalize well to both astrocytic and non-astrocytic peak regions. Then we applied two rounds of fine-tuning. In the first round, we fine-tuned the model using the human BG astrocyte-derived peak universe. We further restricted this set to peaks with variable accessibility across astrocyte subgroups, identified by *crested.pp.filter_regions_on_specificity()* function, to improve model performance on astrocyte-specific peaks (46,722 peaks). We lowered the learning rate to 1e-4 for this step. In the second fine-tuning round, we used a set of highly contrastive differentially accessible regions (DARs; 11,998 peaks) obtained as the union of pairwise subgroup comparisons performed with the *snapatac2.tl.diff_test()* function with a lowered learning rate of 1e-5, to further enhance the model’s ability to discriminate among astrocyte subgroups.

To evaluate the model performance, we used the default CREsted evaluation functions, *crested.pl.scatter.class_density()* and *crested.pl.heatmap.correlations_predictions()*, on the contrastive DARs test set.

We additionally trained a model on the three GM STR clusters (DeepHumanAstroSTR). Here, we followed an identical procedure to the subgroup model training with a first training step on the whole human BG universe, a second fine-tuning step on variable GM STR peaks (119,773 peaks, identified by *crested.pp.filter_regions_on_specificity()*), and a final fine-tuning step on a set of highly contrastive DARs (13,171 peaks) obtained as the union of pairwise subgroup comparisons performed with the *snapatac2.tl.diff_test()* function.

#### CREsted Motif Analysis

To obtain the characteristic motifs per subgroup with the CREsted model, we calculated nucleotide contribution scores for a set of characteristic sequences. For each subgroup, we first defined the top 500 most specific sequences from the set of contrastive DARs based on the averaged specificity of model predictions and peak heights with *crested.pp.utils._calc_proportion()* function. These regions were then combined across subgroups into a unified set of 1,500 regions, for which we calculated contribution scores (method=’integrated_grad’) for each model class. Using tfmodisco-lite (https://github.com/jmschrei/tfmodisco-lite), we extracted the most recurrent motifs that the model considered important for predicting subgroup-specific chromatin accessibility. This analysis allowed us to identify and compare motif importance profiles across astrocyte subgroups.

To infer TF candidates for each motif, we first matched the identified motifs to a reference motif database (*crested.get_motif_db()*).^60^ The resulting matches were then manually curated using marker TFs identified from the snRNA-seq analysis to select the best TF candidates.

To obtain motif comparisons from the DeepMouseBrain3^33^ and DeepHumanBrain^33^ model, we followed the exact same procedure as described above, and calculated contribution scores for relevant classes (319_Astro-TE_NN, 318_Astro-NT_NN, 317_Astro-CB_NN, and 316_Bergmann_NN in DeepMouseBrain3 and ASCT_1, ASCT_2 and ASCT_3 in DeepHumanBrain). The models were loaded using *crested.get_model()*. We then used the motif locations obtained from the original modisco calculations to retrieve the importance scores for each of these classes. For matched TF expression to patterns from DeepMouseBrain3, we used the Yao et al. whole mouse brain dataset.^6^

#### *WIF1* and *SPARC* Gene Locus Analysis

To visualize the gene loci of *WIF1* and *SPARC*, we used pyGenomeTracks.^63^ The coordinates were chr12:64,989,276-65,125,214 (hg38) for *WIF1* and chr5:151,656,390-151,688,948 (hg38) for *SPARC*. For both loci, HiC values were log-transformed, with the display range set from 5 to 16.

#### Tissue Preparation

A separate donor (52-year-old, Male) was obtained from the UW BRaiN Bank for in-situ hybridization (ISH) experiments. For the present study, brain slabs were identified and dissected from anterior and posterior BG, which contained dorsal and ventral striatum, and Pu and GP, respectively. Dissected tissues were then mounted onto a cryosection sample holder and sectioned at -20°C at 16µM intervals. Subsequent sections were then mounted onto standard microscope slides and stored at -80°C until needed for ISH experiments.

#### Combined RNAScope ISH and Immunohistochemistry

Two tissue sections were removed from storage and fixed for 15 minutes in a bath of 4% PFA. After rinsing, sections were then dehydrated in a series of room-temperature ethanol (EtOH) baths of increasing concentrations (50%, 70%, and 2x 100% EtOH), for five minutes each. Following this, RNAScope probes (ACDBio) for AQP4 were applied to each section and were combined with probes against either GRM3 (striatal astrocytes) or SLC6A11 (extra-striatal astrocytes). The multiplex fluorescent RNAscope1 kit (ACD Bio #323100) was used according to the manufacturer’s protocol.^64^ Upon completion of RNAScope probe application, immunohistochemistry (IHC) was performed on all sections for GFAP expression. Sections were blocked in a 1:8 solution of normal goat serum (Jackson Immuno) and 1X PBS for 15 minutes. The primary GFAP antibody was diluted to a concentration of 1:500 with the blocking solution mentioned above. Then, 50µL of the primary was applied to each section and incubated for 24 hours. After this, Alexa Fluor 488 (ThermoFisher Scientific) was diluted to a concentration of 1:500 in blocking solution and 50µL was applied to the slides. The secondary incubation lasted one hour, after which slides were washed, coverslipped, and imaged using a slide scanning imaging microscope (Olympus VS200).

