## Supplemental_Figures_BG_Astrocyte for "Circuit specific specialization of human basal ganglia astrocytes"

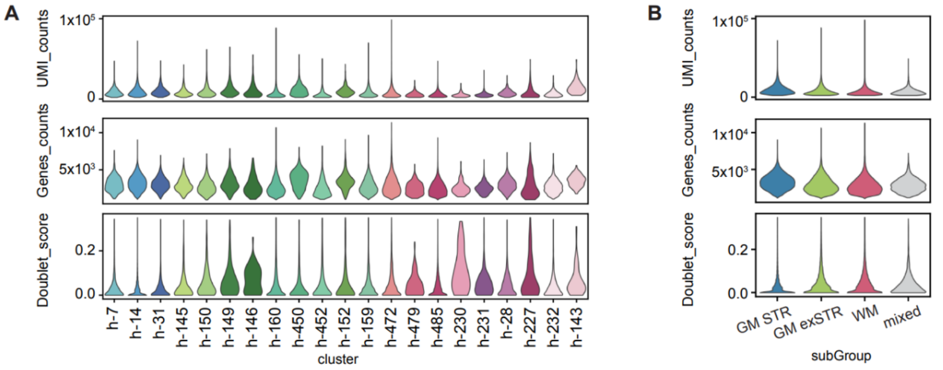


C

| cluster | h-7 | h-14 | h-31 | h-145 | h-150 | h-149 | h-146 | h-160 | h-450 | h-452 | h-152 | h-159 |
|---|---|---|---|---|---|---|---|---|---|---|---|---|
| # of cells | 1,893 | 58,596 | 3,350 | 1,783 | 1,359 | 2,649 | 172 | 17,037 | 1,643 | 2,522 | 1,322 | 12,053 |
| cluster | h-472 | h-479 | h-485 | h-230 | h-231 | h-28 | h-227 | h-232 | h-143 | | | |
| # of cells | 12,864 | 243 | 12,908 | 78 | 1,998 | 1,492 | 162 | 2,469 | 91 | | | |

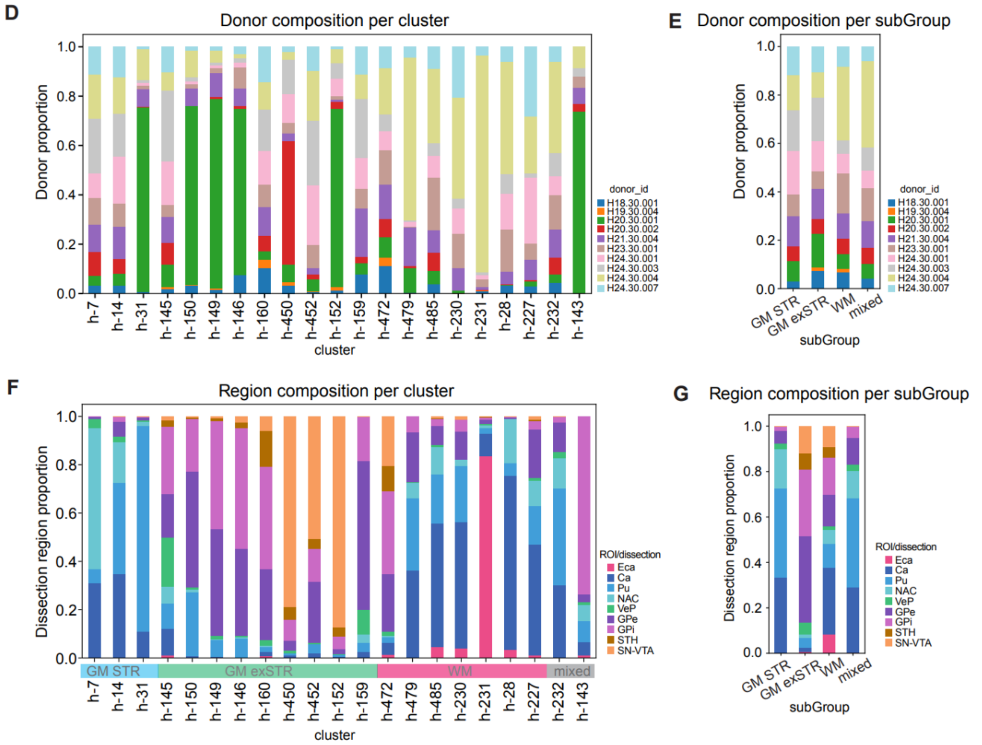


**Figure S1. Quality control and sample composition of human snRNA-seq data, related to Figure 1.**
(**A**) Violin plots showing median UMI counts, detected genes, and doublet scores per cluster.
(**B**) Violin plots showing median UMI counts, detected genes, and doublet scores per subgroup.
(**C**) Table summarizing cell counts per cluster.
(**D**) Bar plot showing donor proportions across clusters. Donor H20.30.001 is slightly enriched in several clusters, consistent with this donor's comparatively higher expression of stress-responsive transcripts (e.g., *EGR1*).
(**E**) Donor composition at the subgroup level.
(**F**) Regional composition of each cluster. Cluster h-231 is predominantly derived from the Eca region (donor H24.30.004; panel D), consistent with this donor being the major source of Eca tissue and further supported by its spatial localization (Figure S2).
(**G**) Regional composition of each subgroup.

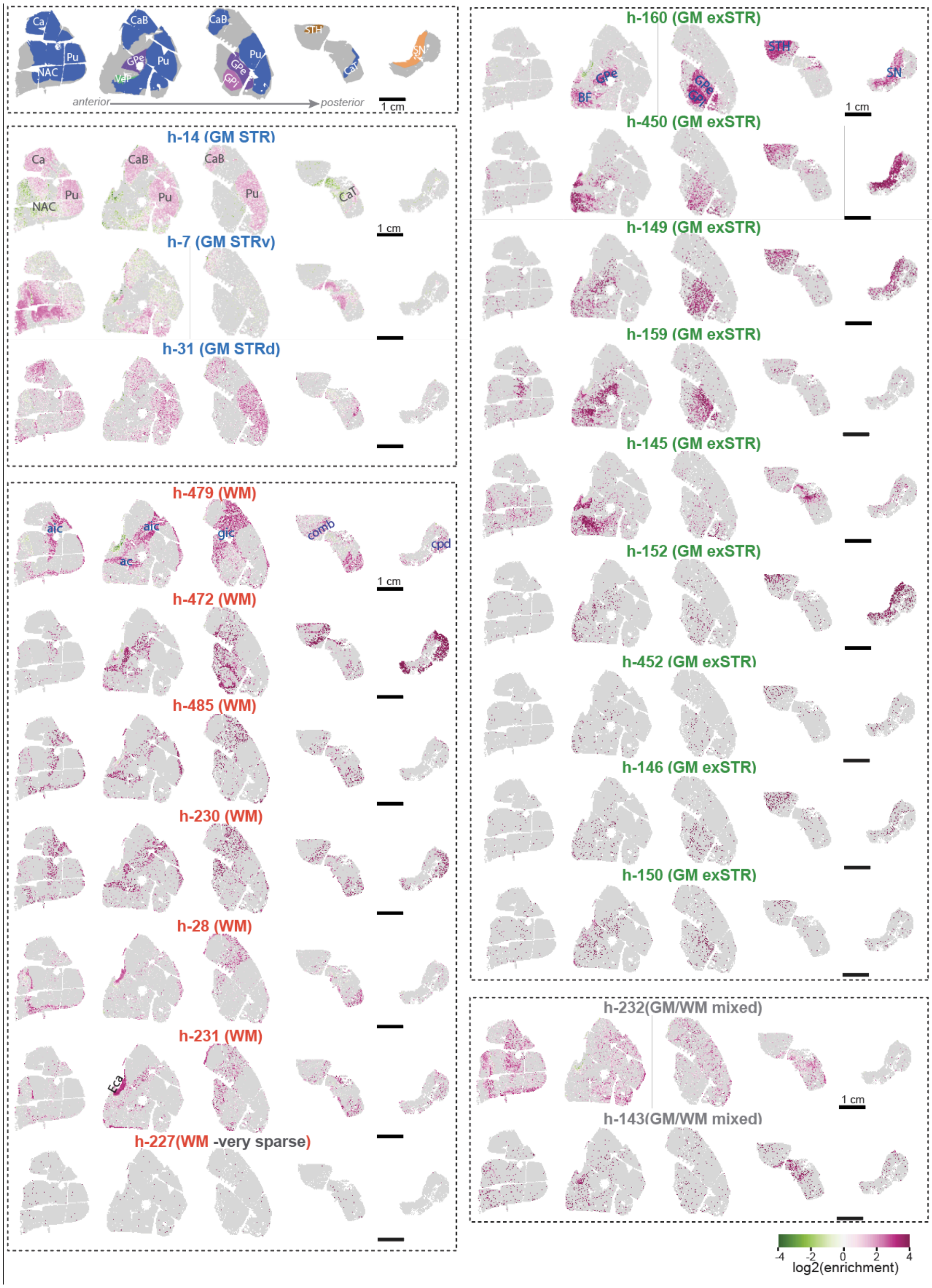


**Figure S2. Spatial enrichment maps of human BG astrocyte clusters, related to Figure 1.**
Spatial enrichment maps of 21 astrocyte clusters across the human BG tissue sections. Dashed rectangles indicate anatomical structures and subgroup categories. Color scale shows $\log_2$(enrichment), calculated as $\log_2$ of the ratio between the proportion of cells from the subgroup within each 300-μm grid cell (“local proportion”) and the proportion of cells from the same subgroup across that section (“global proportion”).

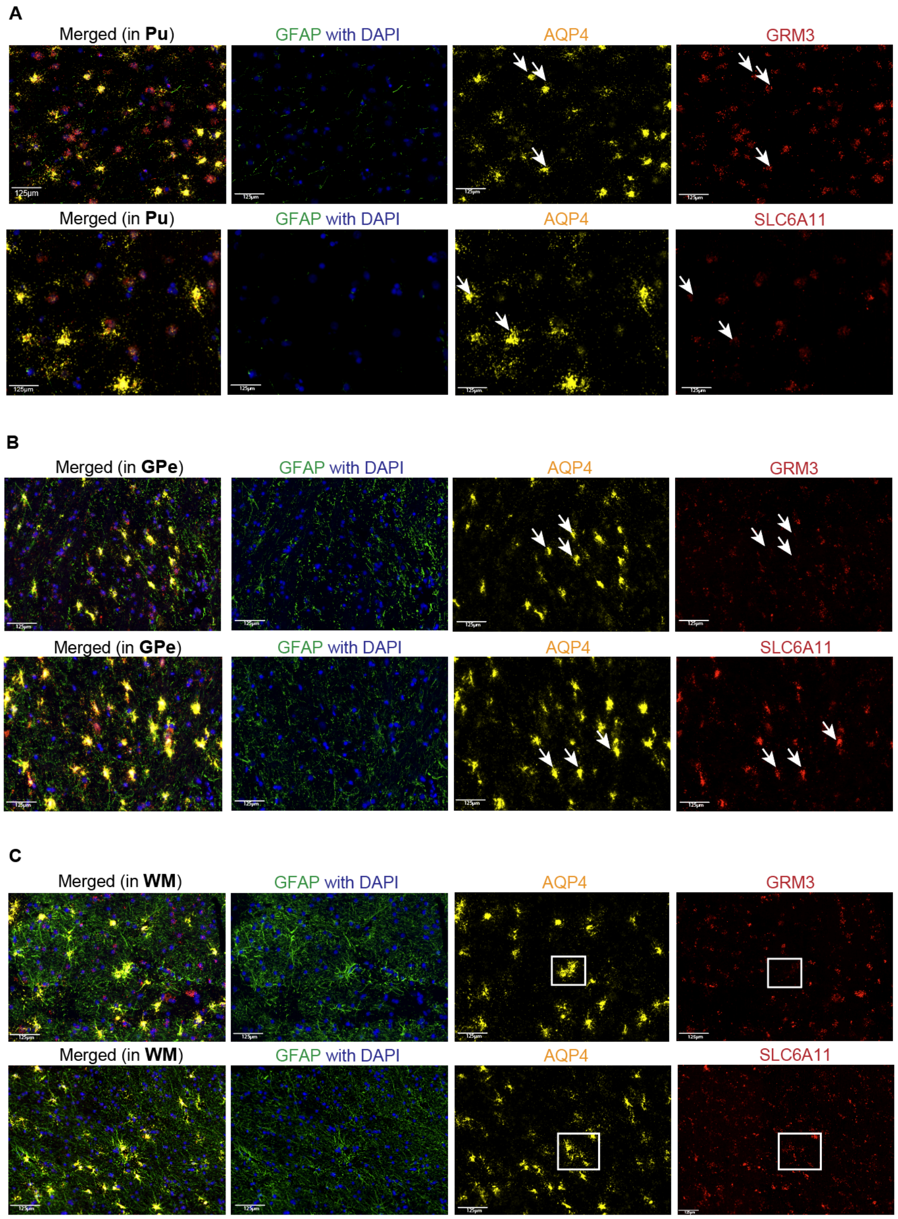


**Figure S3. Distinct astrocyte marker expression and morphology across GM STR, GM exSTR, and WM, related to Figure 1.**

(**A**–**C**) Joint IHC (GFAP) and RNAscope labeling for AQP4 together with either GRM3 or SLC6A11 in the Pu (panel A, representing the GM STR region), the GPe (panel B, representing the extra-striatal GM region), and WM (panel C, representing the WM region).

Across all regions, AQP4 robustly marks astrocytes, while GFAP highlights region-specific morphological organization. WM astrocytes display dense, aligned fibrous processes; GPe (GM exSTR) exhibits an intermediate meshwork of GFAP$^+$ fibers, and Pu (GM STR) astrocytes show comparatively lower GFAP fiber density.

RNAscope signals reveal region-dependent molecular specialization. In the Pu (A), AQP4$^+$ astrocytes prominently express GRM3 (arrows), aligning with the snRNA-seq defined GM STR astrocyte molecular signature. In contrast, SLC6A11 labeling is limited in Pu but becomes enriched in GPe (B) and WM (C), matching the transcriptomic shift toward GABA transporter high astrocyte states characteristic of GM exSTR and WM populations. Insets and arrows highlight representative transcript-positive astrocytes. Scale bars: 125μm.

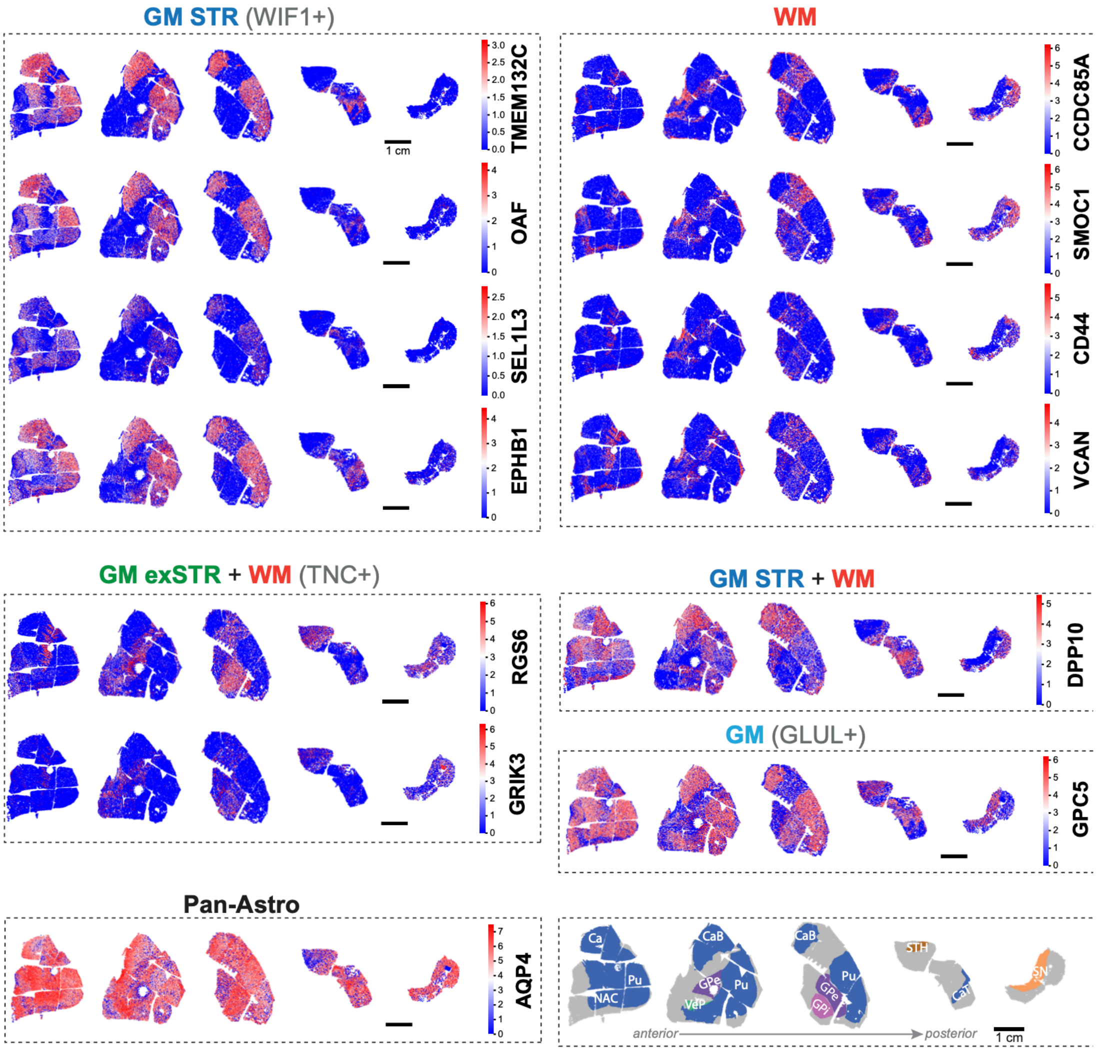


**Figure S4. Spatial expression maps of representative genes, related to Figure 1.**
Spatial expression maps of representative genes from the human BG MERSCOPE panel, shown for astrocytes only, demonstrating that their spatial patterns are consistent with the defined astrocyte subgroup distributions in Figure 1.

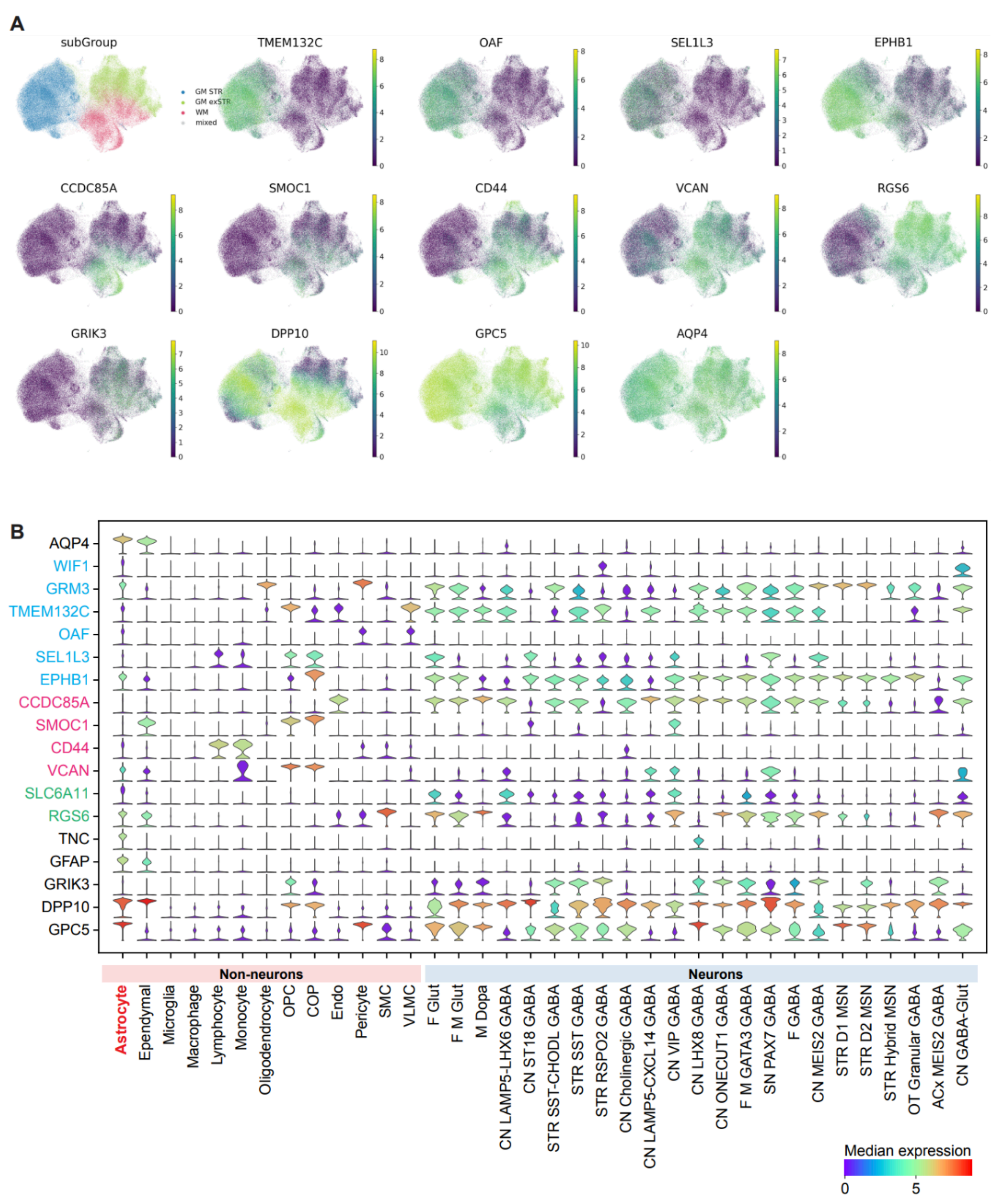


**Figure S5. Representative gene expression patterns of human snRNA-seq data, related to Figure 1.**
(**A**) Expression patterns of the selected genes across human BG astrocytes in the snRNA-seq dataset, corresponding to genes showing clear spatial patterns in Figure S3.
(**B**) Expression patterns of the same gene list, together with those in Figure 1D, across the human BG taxonomy (Subclass level).

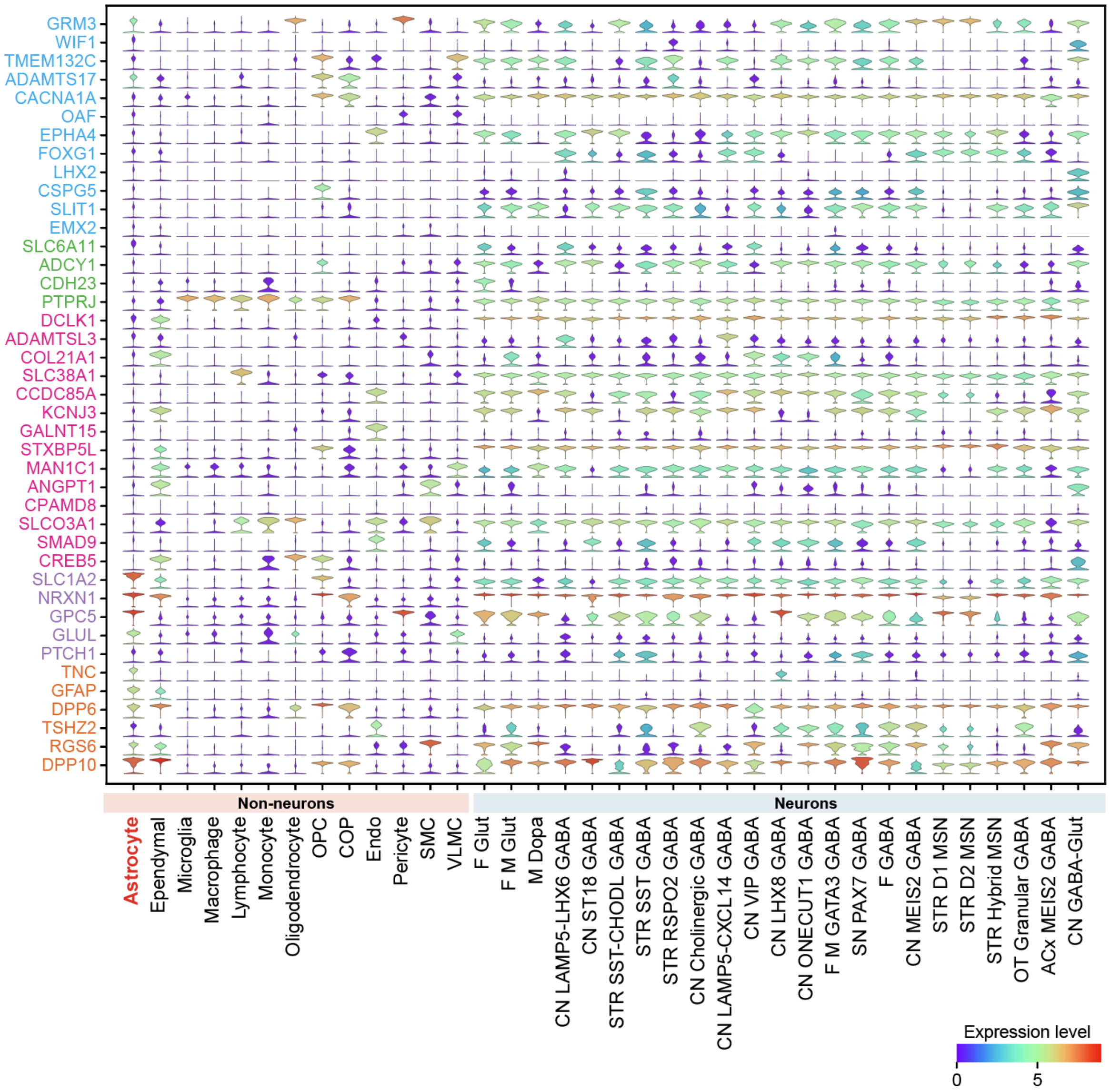


**Figure S6. Violin plots of top DEGs from Figure 2B across the human BG taxonomy at the Subclass level in human snRNA-seq data, related to Figure 2.**

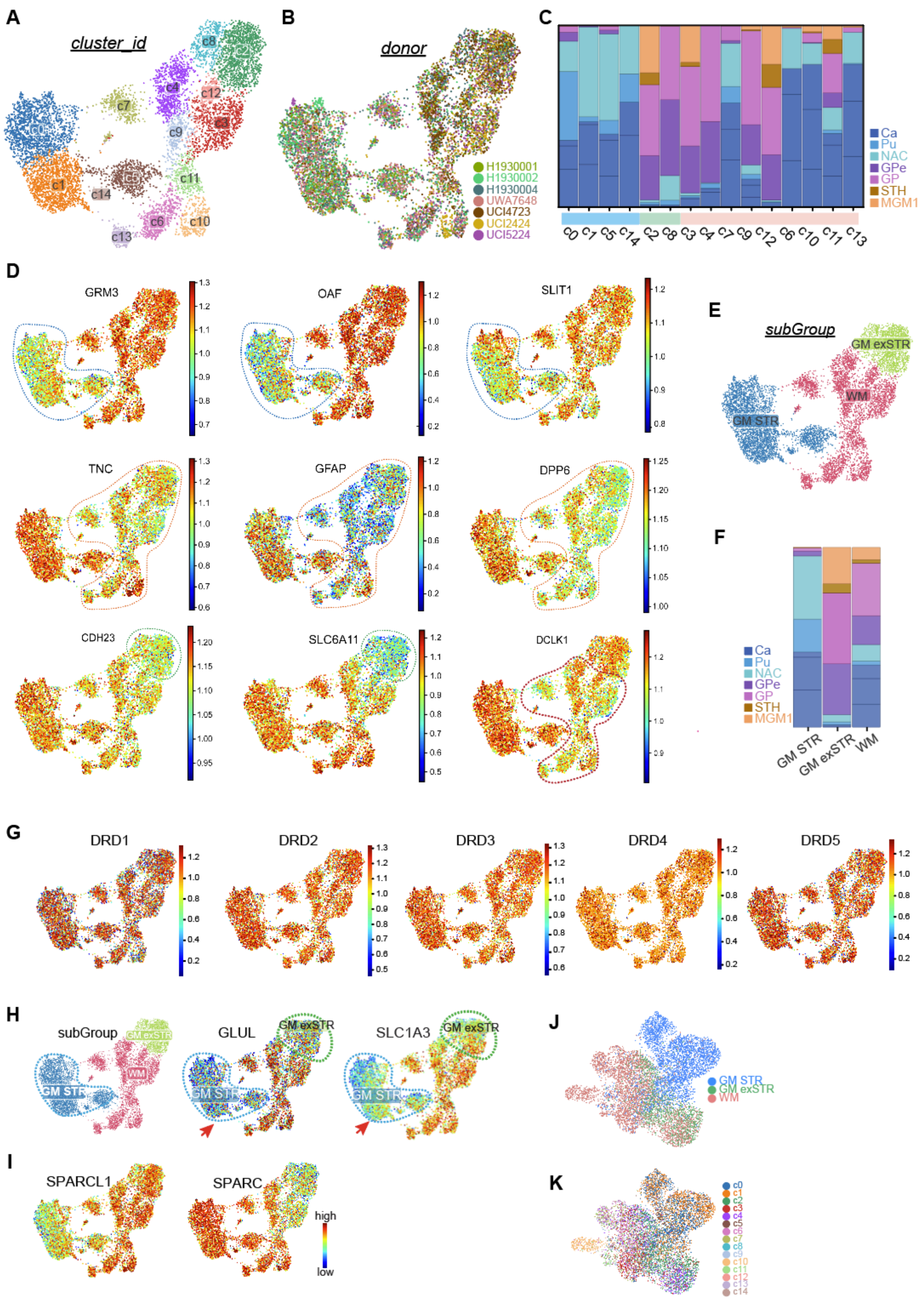


**Figure S7. DNA methylation-based identification and molecular characterization of astrocyte subgroups in the human BG snm3C-seq data, related to Figure 2.**

(**A**-**B**) UMAP plots of mCG profiles from human BG (snm3C-seq). (A) Points colored by cluster_id. (B) Points colored by donor.

(**C**) Regional composition of each cluster. MGM1, a mixture of ventral tegmental region of midbrain (VTR), SN, and red nucleus (RN).
(**D**) UMAP plots showing mCG levels of representative subgroup-specific marker genes identified in Figure 2. GM STR astrocyte markers: *GRM3*, *OAF*, and *SLIT1*; markers shared by GM exSTR and WM: *TNC*, *GFAP*, and *DPP6*; GM exSTR markers: *CDH23* and *SLC6A11*; WM marker: *DCLK1*.
(**E**) UMAP plot of human BG astrocyte methylation profiles, colored by subgroup.
(**F**) Regional composition of each subgroup.
(**G**) UMAP plots showing mCG levels of dopamine receptor genes.
(**H**) Methylation profiles of *GLUL* and *SLC1A3*.
(**I**) Methylation profiles of *SPARCL1* and *SPARC*.
(**J**-**K**) UMAP plots based on 100-kb-resolution 3D chromatin interaction profiles, colored by methylation-derived subgroup annotation (J) and cluster_id (K).

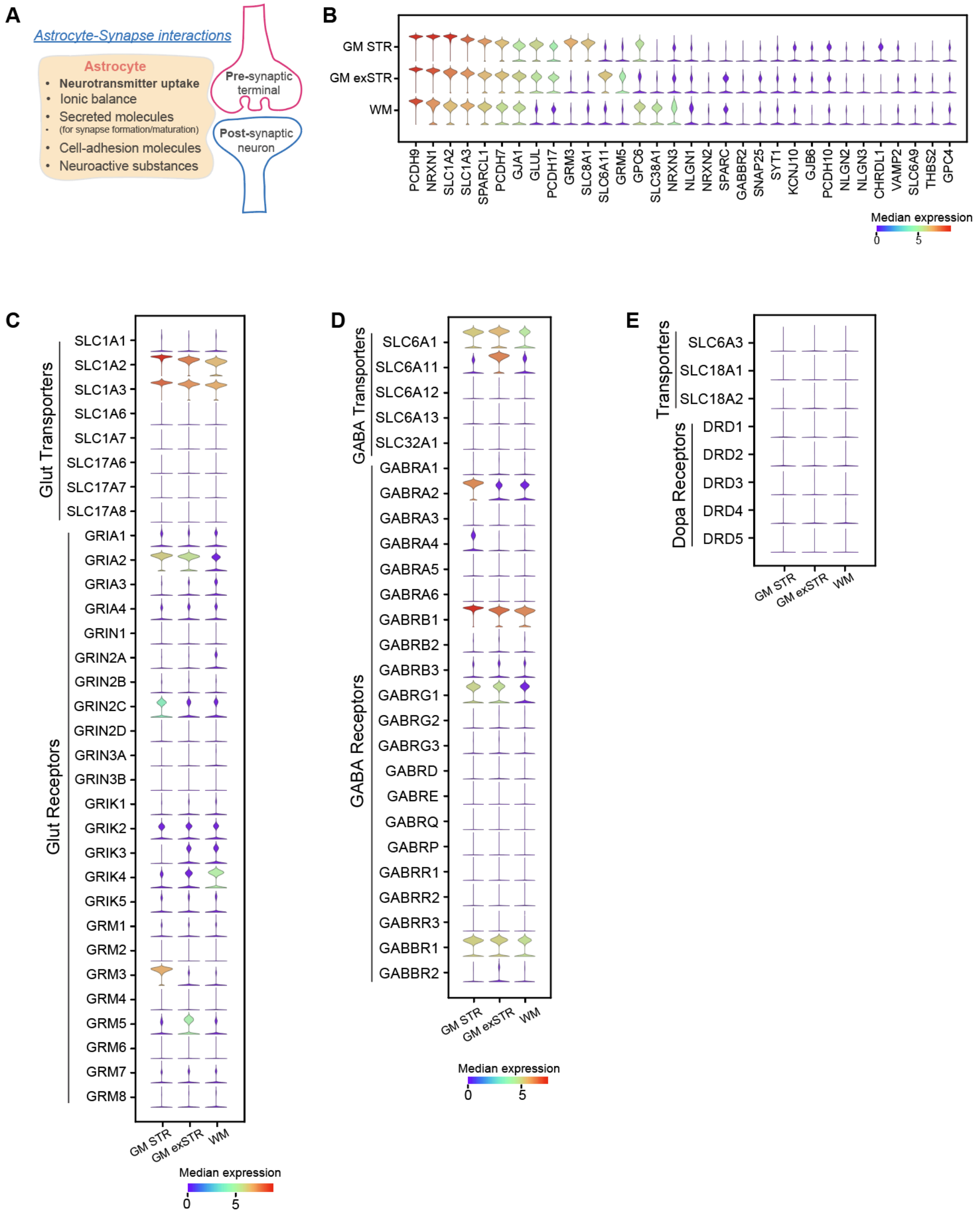


**Figure S8. Expression patterns of genes associated with astrocyte-synapse interactions in the human snRNA-seq data, related to Figure 2.**
(**A**) Schematic illustrating astrocyte-synapse interactions.
(**B**) Expression patterns of core genes involved in the interactions shown in (A).
(**C**) Expression patterns of glutamate receptor and transporter genes across astrocyte subgroups.
(**D**) Expression patterns of GABA receptor and transporter genes across astrocyte subgroups.
(**E**) Expression patterns of dopamine receptor and transporter genes across astrocyte subgroups.

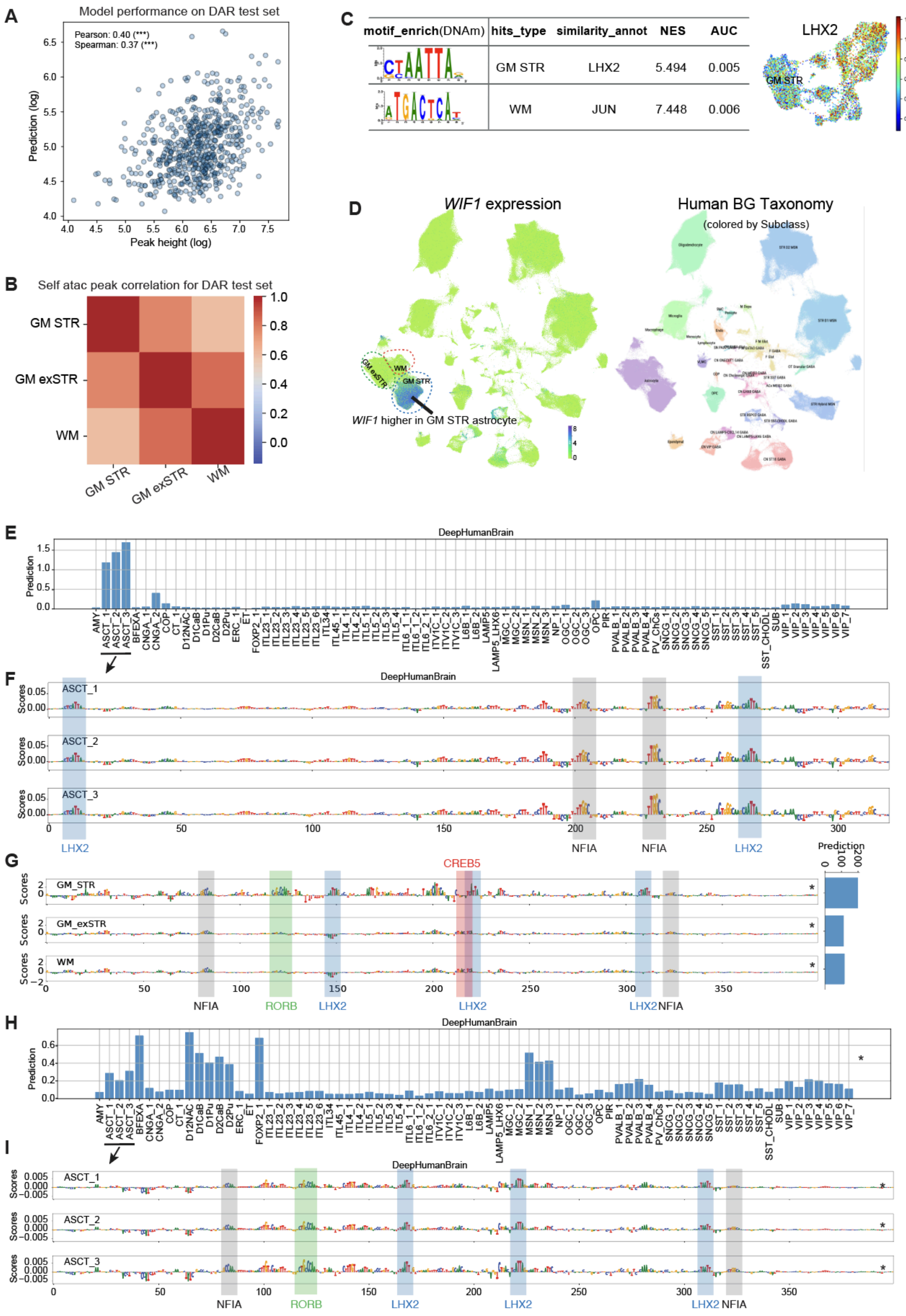


**Figure S9. Multimodal validation of the BG astrocyte CREsted model predictions using independent model outputs and snm3C-seq methylation data, related to Figure 3.**

(**A**) Scatter plot of log-transformed CREsted model predictions versus peak heights on a set of held out test set regions from the contrasting DAR set. ***, P-value < 0.001 (P-values from two-sided hypothesis tests).

(**B**) Heatmap of self-correlation of log-transformed peak heights across subgroups, indicating inter-subgroup similarity of accessibility.
(**C**) Enrichment analysis of subgroup-specific DMRs from the human snm3C-seq data showing LHX2 motif enrichment in GM STR astrocytes, and JUN motif enrichment in WM, consistent with model predictions. Right, LHX2 hypomethylation in GM STR astrocytes.
(**D**) *WIF1* showed specific high expression in GM STR astrocytes across the human BG taxonomy in the snRNA-seq data. Left: *WIF1* expression levels with outlines marking astrocyte subgroups. Right: human BG taxonomy colored by Group.
(**E**) DeepHumanBrain (Hecker et al., 2025) CREsted model predictions on the identified *WIF1* enhancer candidate.
(**F**) DeepHumanBrain model contribution scores on the identified *WIF1* enhancer candidate for the three telencephalic astrocyte groups.
(**G**) CREsted model contribution scores across astrocyte subgroups for the second *WIF1* enhancer candidate (indicated by an asterisk in Figure 3D).
(**H**) DeepHumanBrain CREsted model predictions on the second identified *WIF1* enhancer candidate (indicated by an asterisk in Figure 3D).
(**I**) DeepHumanBrain model contribution scores across the three telencephalic astrocyte groups for the second identified *WIF1* enhancer candidate (indicated by an asterisk in Figure 3D).

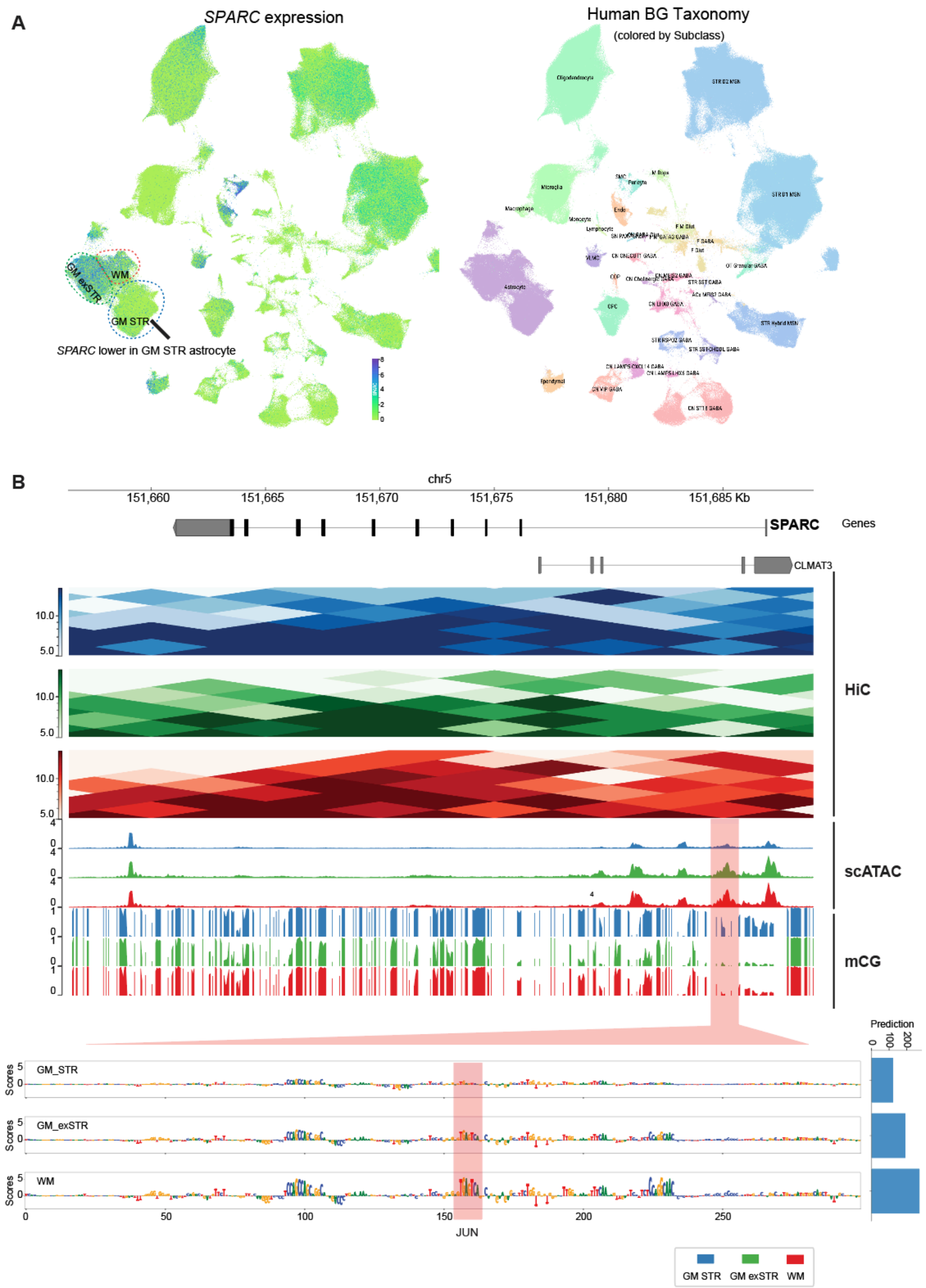


**Figure S10. Multimodal characterization of *SPARC* expression and regulatory features across human BG astrocyte subgroups, related to Figure 3.**

(**A**) *SPARC* shows higher expression in GM exSTR and WM astrocytes across the human BG taxonomy in the snRNA-seq data. Left: *SPARC* expression levels with outlines marking astrocyte subgroups. Right: human BG taxonomy colored by Group.

(**B**) Multi-omic view of the *SPARC* gene locus, with Hi-C contact maps, and snATAC-seq and mCG tracks. The CREsted model highlights a WM/GM exSTR-specific regulatory region inside the *SPARC* gene locus predicts this region to be selectively accessible in WM/GM exSTR astrocytes.

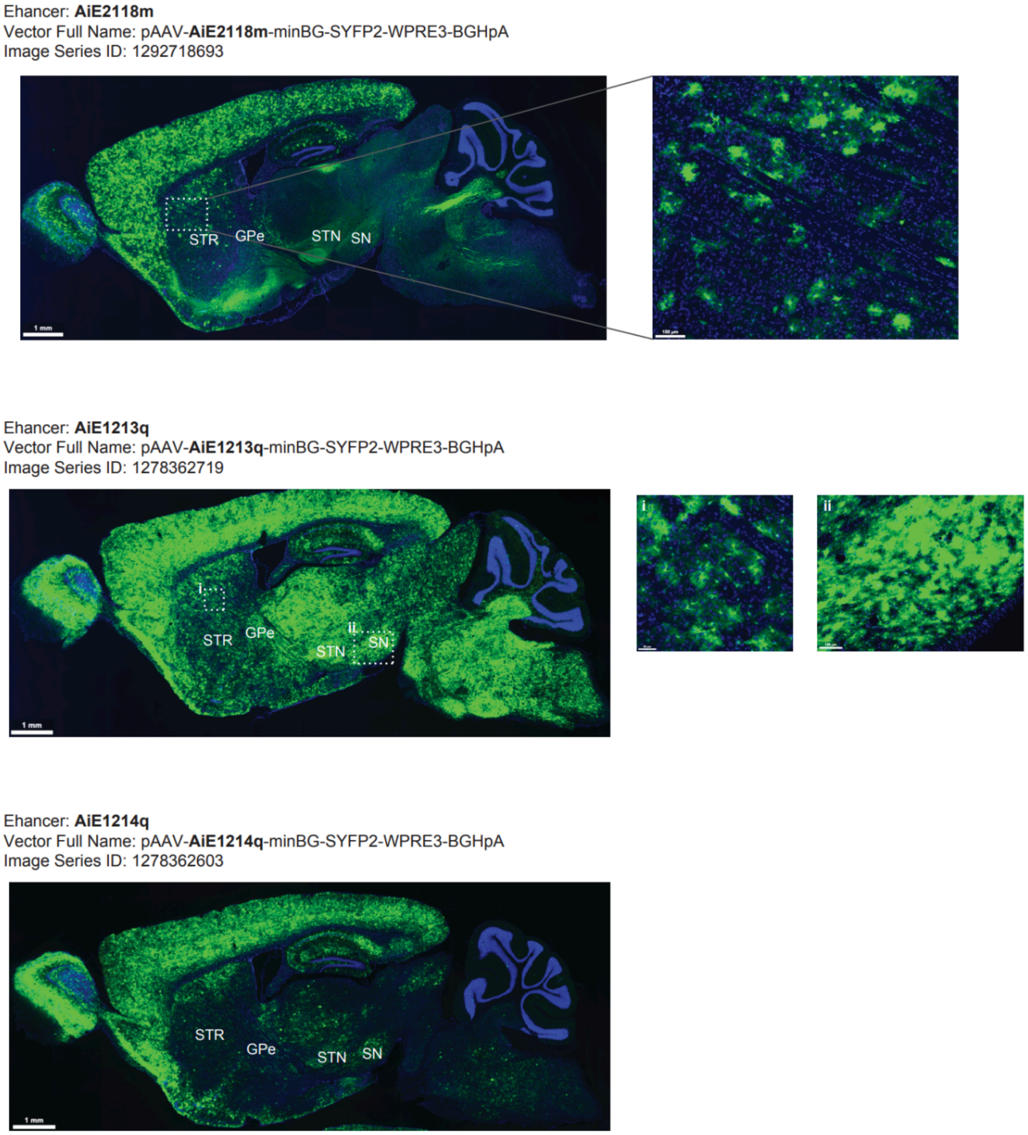


**Figure S11. Experimental validation of model-predicted enhancer activity using viral labeling data from the Allen Brain Map Genetic Tools Atlas, related to Figure 4.**
(**A-C**) Epifluorescence images showing in vivo viral labeling patterns for three enhancer constructs corresponding to model-predicted regulatory regions in Figure 4A: AiE2118m (A), AiE1213q (B), and AiE1214q (C). Images were obtained from the Allen Institute Genetic Tools Atlas (https://brain-map.org/bkp/experiment/genetic-tools/genetic-tools-atlas). SYFP2 (green) reporting enhancer-driven expression and DAPI (blue) labeling nuclei.

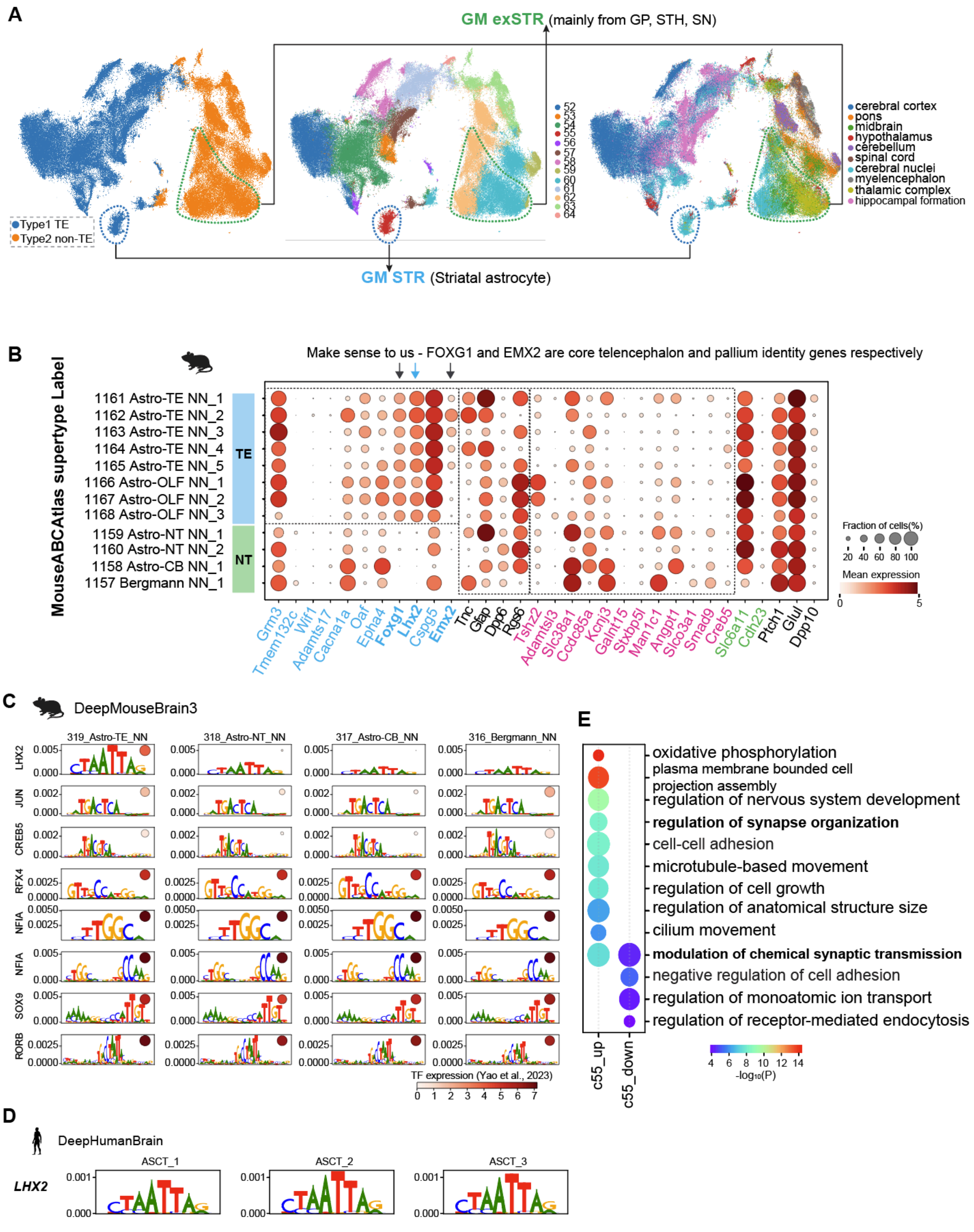


**Figure S12. Cross-species and cross-modal evidence showing transcriptional and regulatory programs in astrocytes, related to Figure 5.**

(**A**) UMAP plots of human whole-brain snRNA-seq astrocyte data (Siletti et al., 2023), colored by Type (TE = telencephalon, non-TE = non-telencephalon), by cluster, and by dissection tissue.

(**B**) Dot plot showing the expression patterns of the same gene list as in Figure 5B across astrocyte clusters in the mouse whole-brain dataset (Yao et al., 2023). Although many genes differed between human and mouse, *FOXG1*, *LHX2*, and *EMX2* showed conserved expression patterns in mouse. TE, telencephalon; OLF, olfactory; NT, non-telencephalon; CB, cerebellum; NN, non-neuronal cells.
(**C**) Motifs identified by the DeepMouseBrain3 (Hecker et al., 2025) model in the top 500 most specifically accessible regions for each astrocyte subgroup. Motif heights are the average contribution scores from all instances identified across all mouse astrocyte classes. The distribution of motif instances found in each subgroup region set is shown with bar plots on the right. The absolute TF expression (logCPM) of proposed motif-TF matches is shown in color. Relative TF expression (per row divided by maximum value) is indicated by dot size.
(**D**) Same as (C), but with the DeepHumanBrain (Hecker et al., 2025) model and only focusing on the LHX2 motif. Contribution scores were calculated for the three human telencephalic astrocyte classes. ASC, astrocyte.
(**E**) GO enrichment analysis of upregulated and downregulated genes identified in the c55 striatal astrocyte cluster shown in Figure 5D. Each dot represents an enriched biological process. Dot color, enrichment significance (-$\log_{10}$ P value). Dot size, gene set size of GO terms.

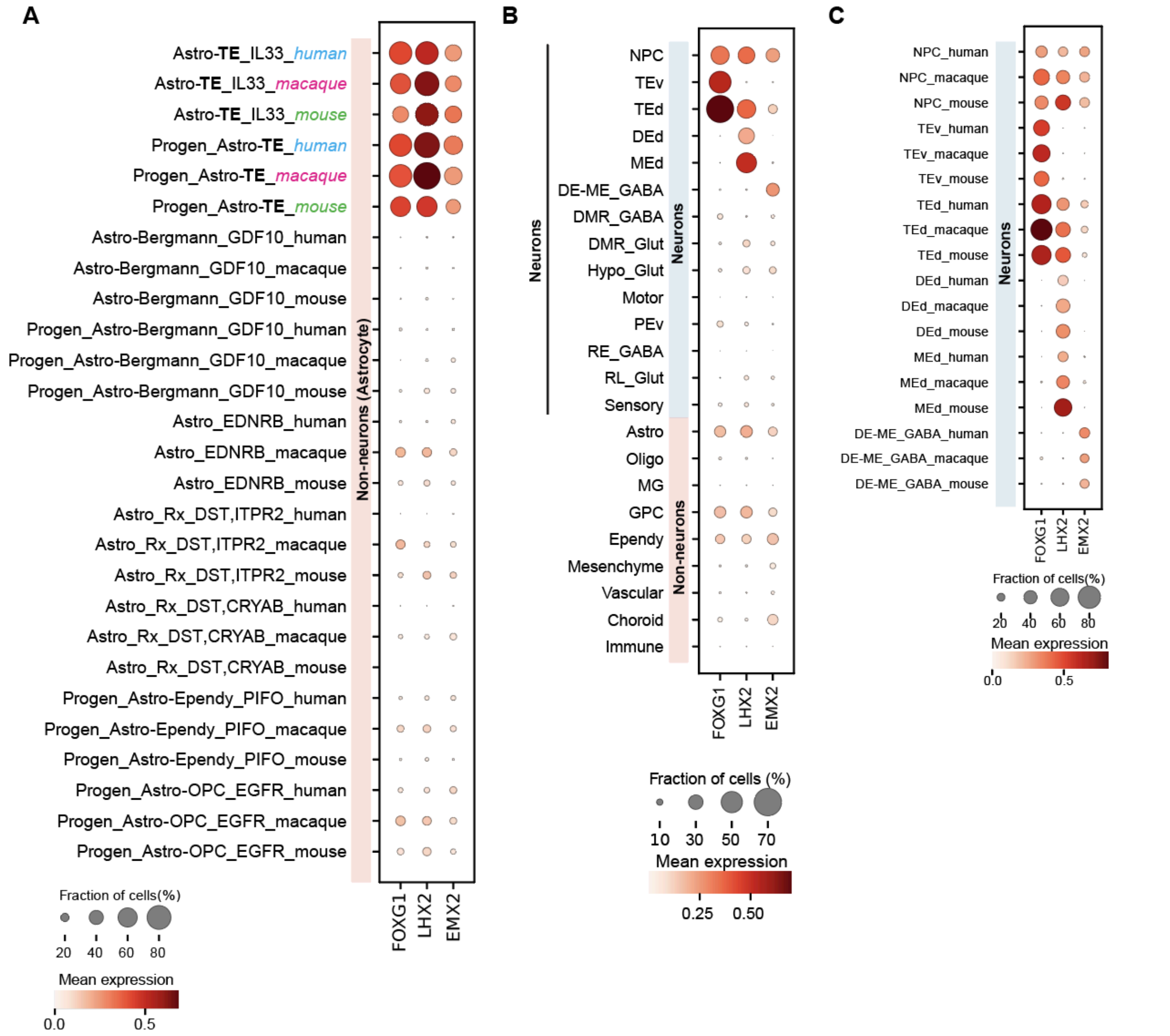


**Figure S13. Species-conserved but lineage-divergent expression patterns of *LHX2*, *FOXG1*, and *EMX2* in whole-brain developmental datasets (Schmitz et al., 2025), related to Figure 5.**

(**A**) Dot plot showing that *LHX2* is highly expressed in telencephalic (TE) astrocytes and their progenitors across prenatal human, macaque, and mouse datasets (Schmitz et al., 2025), resembling the patterns of *FOXG1* and *EMX2*. Astro_Rx_DST, *DST*$^{+}$ reactive-like astrocytes.

(**B**) Dot plots showing divergent expression patterns of *FOXG1*, *LHX2*, and *EMX2* across neurons in the aggregated prenatal datasets from human, macaque, and mouse (same datasets from Schmitz et al., 2025). All three genes are highly expressed in neuroepithelial/neural progenitor cells (NPCs). *FOXG1* is enriched throughout the telencephalon, including both dorsal (TEd) and ventral (TEv) regions; *LHX2* is preferentially expressed in dorsal domains (DEd, MEd, and TEd); and *EMX2* is mainly enriched in DE-ME GABAergic populations. DE-ME_GABA, diencephalic/mesencephalic GABAergic-like neurons. DEd, dorsal thalamic neurons. MEd, dorsal mesencephalic neurons. DMR_GABA, diencephalic-mesencephalic-rhombencephalic GABAergic neurons. Hypo_Glut, hypothalamic glutamatergic neurons. PEv, secondary prosencephalic neurons. GPC, glial progenitor cells. MG, microglia.

(**C**) Corresponding dot plots showing the same gene expression patterns separated by species (human, macaque, and mouse) for the cell types highlighted in panel (B).

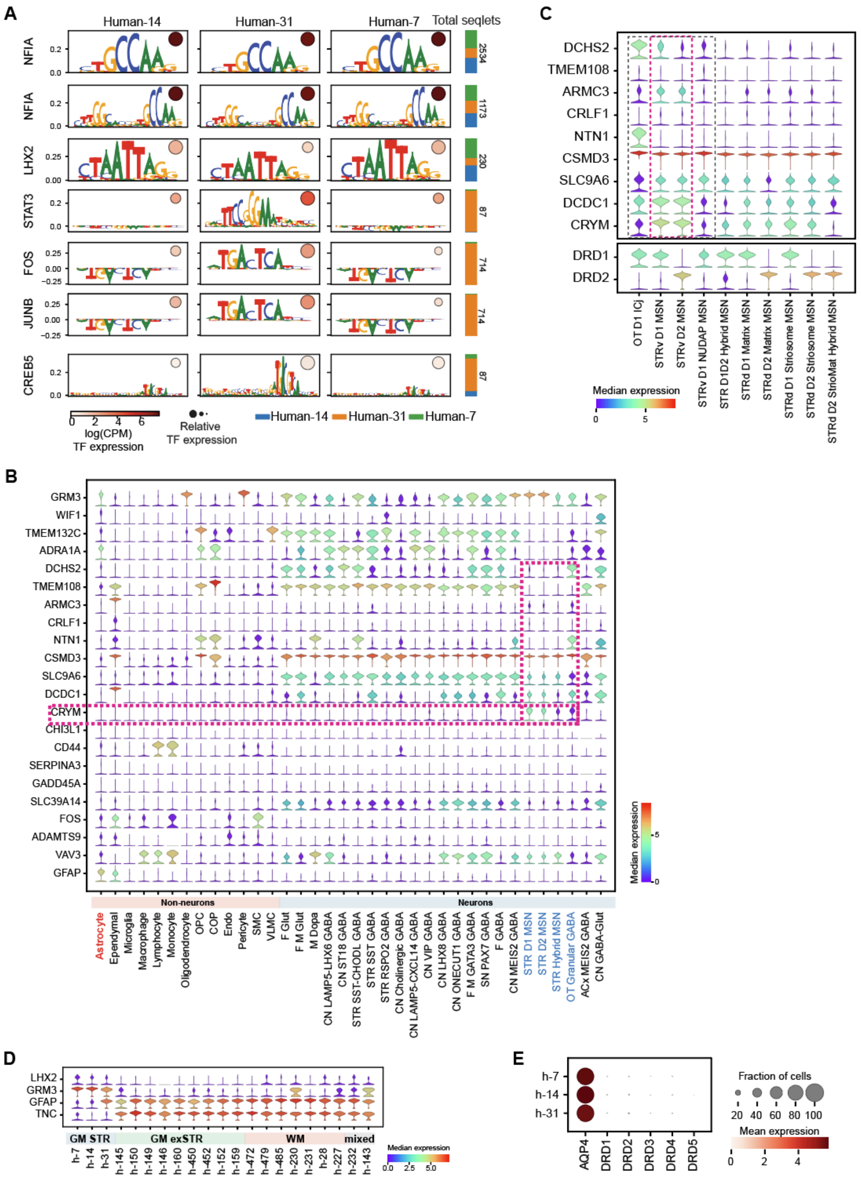

**Figure S14. Cross-taxonomy expression patterns of GM STR astrocyte local heterogeneity markers, related to Figure 6.**

(**A**) Motifs identified by the model in the top 500 most specifically accessible regions for each cluster by the cluster-model. JUN/FOS, Activator Protein-1 components. Motif heights are the average contribution scores from all instances identified across all GM STR astrocyte clusters. The distribution of motif instances found in each cluster region set is shown with bar plots on the right. The absolute TF expression (logCPM) of proposed motif-TF matches is shown in color. Relative TF expression (per row divided by maximum value) is indicated by dot size.

(**B**) Violin plots showing the expression of top DEGs identified in Figure 6B across the human BG taxonomy at Subclass level in the human BG snRNA-seq data.

(**C**) Violin plots showing the expression of top DEGs identified in Figure 6B across the human BG MSNs at Group level in the human BG snRNA-seq data.

(**D**) Violin plots showing the expression patterns of representative BG astrocyte subgroup marker genes across clusters in the human BG snRNA-seq data.

(**E**) Dot plot showing the expression of AQP4 and dopamine receptor genes in the human BG snRNA-seq data.

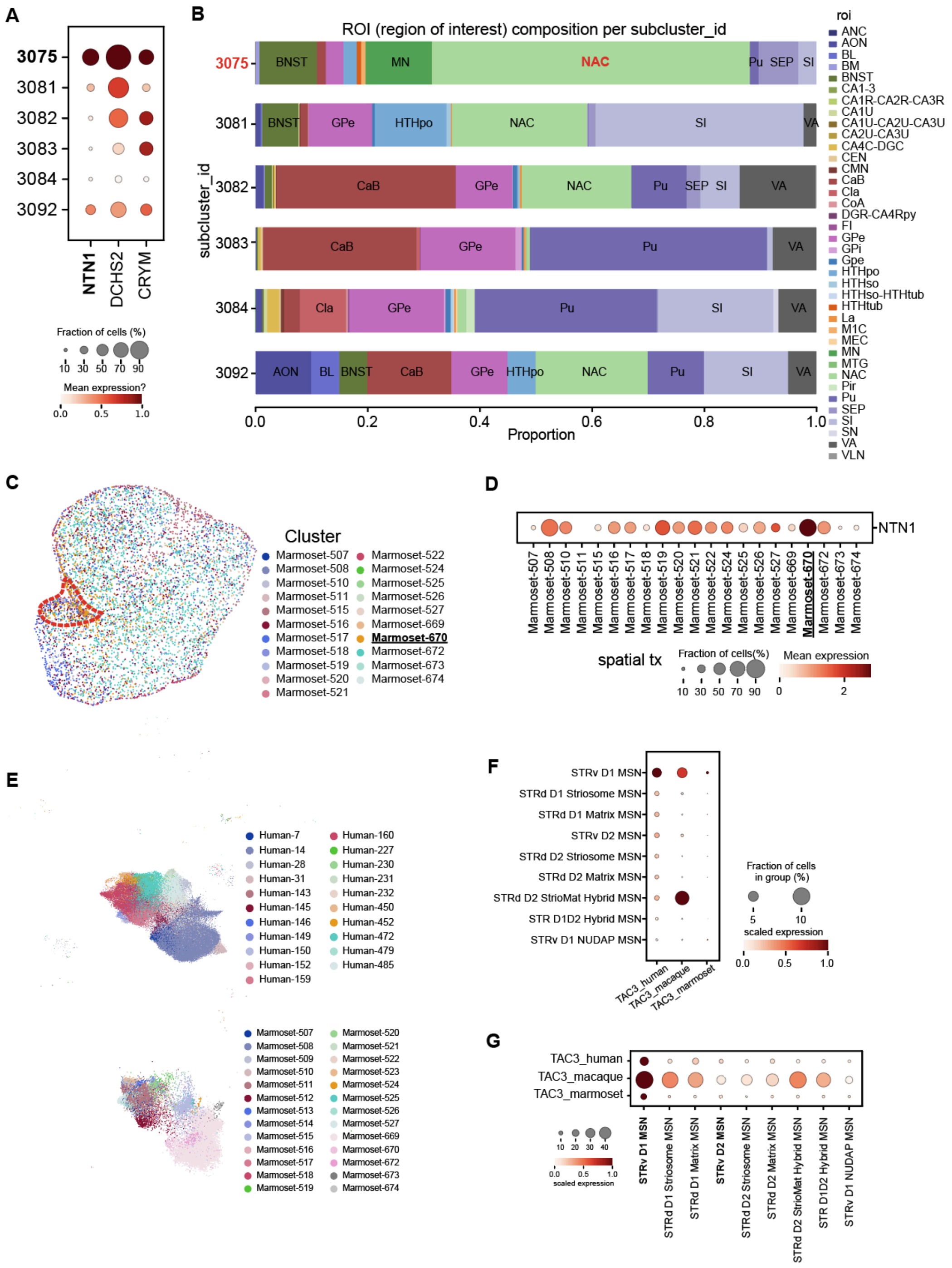


**Figure S15. Cell populations enriched in the ventral striatum, related to Figure 6.**
(**A**) Expression patterns of h-7 marker genes across subclusters within cluster 55 (striatal astrocytes) of the Siletti et al. whole-brain atlas. Color bar indicates scaled expression values normalized by gene.
(**B**) Regional composition of subclusters within the cluster 55. The subcluster_id 3075, which shows the highest expression levels in (A), is primarily sampled from the NAC region, in line with our Figure 6 results.
(**C**) Spatial transcriptomic map of marmoset clusters in the Astrocyte Group, generated from the companion paper by Hewitt et al. The dotted outline denotes the primary distribution of the Marmoset-670 cluster, which largely overlaps with the NACsmd.

(**D**) *NTN1* expression patterns across marmoset astrocyte clusters using the same spatial datasets in (C).
(**E**) Integrated UMAP of human and marmoset MSN snRNA-seq datasets, colored by clusters. The upper panel shows human cells, and the lower panel shows marmoset cells.
(**F-G**) *TAC3* expression across MSN Groups in human, macaque, and marmoset. (F) shows the snRNA-seq datasets, and (G) shows the spatial data.
